# Differential migration mechanics and immune responses of glioblastoma subtypes

**DOI:** 10.1101/2022.06.26.497270

**Authors:** Ghaidan A. Shamsan, Chao J. Liu, Brooke C. Braman, Ruyi Li, Susan K. Rathe, Aaron L. Sarver, Nima Ghaderi, Mariah M. McMahon, Rebecca L. Klank, Barbara R. Tschida, S. Joey McFarren, Pamela C. Rosato, David Masopust, Jann N. Sarkaria, H. Brent Clark, Steven S. Rosenfeld, David A. Largaespada, David J. Odde

## Abstract

Glioblastoma remains a deadly cancer driven in part by invasion of tumor cells into the brain. Transcriptomic analyses have identified distinct molecular subtypes, but mechanistic differences that account for clinical differences are not clear. Here, we show that, as predicted by the motor-clutch model of cell migration, mesenchymal glioma cells are more spread, generate larger traction forces, and migrate faster in brain tissue compared to proneural cells. Despite their rapid migration and comparable proliferation rates in vitro, mice with mesenchymal tumors survive longer than those with proneural tumors. This improved survival correlated with an immune response in the mesenchymal tumors, including T cell-mediated. Consistently, inducing mesenchymal tumors in immunodeficient mice resulted in shorter survival supporting a protective immune role in mesenchymal tumors. Thus, mesenchymal tumors have aggressive migration, but are immunologically ‘hot’ which suppresses net proliferation. These two features counteract each other and may explain the lack of a strong survival difference between subtypes clinically, while also opening up new opportunities for subtype-specific therapies.

**Significant Statement:** This study highlights new mechanical and immunological insights into glioblastoma molecular subtypes using an integrated modeling-genome engineering strategy, which can potentially facilitate glioblastoma subtype-specific therapeutic strategies.

## INTRODUCTION

Glioblastoma (GBM: WHO grade IV primary brain tumor) progression can be characterized in terms of tumor growth and spreading, two key parameters which are influenced by many of the hallmarks of cancer (Hanahan and Weinberg, 2011). In GBM, tumor spreading is driven by tumor cells’ ability to infiltrate healthy brain parenchyma, which prevents complete surgical resection and results in tumor recurrence (Lefranc, Brotchi and Kiss, 2005; Hoelzinger, Demuth and Berens, 2007; de Gooijer *et al*., 2018). Molecular and genetic analyses of human GBM have identified at least three distinct molecular subtypes: proneural, classical, and mesenchymal (Phillips *et al*., 2006; Verhaak *et al*., 2010; Wang *et al*., 2017). These subtypes were shown to strongly correlate with specific genetic alterations (Mesenchymal: *NF1* loss; Classical: *EGFRvIII*; Proneural: *PDGFRA*) and cellular developmental states (Verhaak *et al*., 2010; Patel *et al*., 2014; Wang *et al*., 2017; Neftel *et al*., 2019). Despite accumulating evidence of distinct transcriptomic and genetic signatures, the characteristic mechanistic differences between such signatures, if any, have not been identified. As a result, it remains unclear how knowledge of the different subtypes should inform clinical decisions.

One intriguing correlate of subtype is the level of CD44 expression, a cell surface protein expressed on tumor and immune cells, which is known to play a role in cancer progression across a variety of cancers including GBM (Naor *et al*., 2002; Toole, 2009; Bhat *et al*., 2013; Mao *et al*., 2013; Ozawa *et al*., 2014; Pietras *et al*., 2014; Mooney *et al*., 2016; Klank *et al*., 2017a; Wang *et al*., 2017; Neftel *et al*., 2019). In GBM, we previously showed that CD44 expression is a prognostic marker with a biphasic dependence: better outcomes are observed at both lower and higher levels of CD44 while poorer outcomes are observed at intermediate levels, an example of optimality and the ‘goldilocks’ phenomenon (Klank *et al*., 2017a). In animal models, CD44 expression has further been shown to correlate with glioma cell migration in a biphasic relationship with a peak migration rate at intermediate expression level, which also correlated with the minimum in survival in both the animal model and human GBM (Klank *et al*., 2017a). In addition, *CD44* transcript levels are shown to vary across GBM molecular subtypes with elevated expression in mesenchymal tumors (Phillips *et al*., 2006; Verhaak *et al*., 2010; Pietras *et al*., 2014). *CD44* expression in the mesenchymal tumors is, on average, closer to the *CD44* level that corresponds to the minimum in patient survival than the proneural subtype (Klank *et al*., 2017a). As an adhesion molecule, CD44 engages the extracellular matrix with the actin cytoskeleton through adapter proteins to mediate cell migration (Toole, 2009). This suggests that mesenchymal cells have a near-optimal level of CD44 adhesion molecules to serve as molecular “clutches” that resist myosin II motor forces, allowing them to migrate faster than proneural cells which on average have a lower, suboptimal level of CD44 clutches (see Figure 2C and 4E in Klank *et al.,* 2017). This could then explain the slightly worse outcomes for mesenchymal patients and higher cell migration and invasion (Yoshida *et al*., 2012; Wang *et al*., 2017). In addition, it would predict that mesenchymal cells would have lager cellular spread area, be more polarized, and generate more traction force as they migrate. More generally, lower CD44 is indicative of an epithelial state and higher CD44 indicative of a mesenchymal state (Bloushtain-Qimron *et al*., 2008; Polyak and Weinberg, 2009), and so an increase in myosin motors and adhesions, either integrin- or CD44-mediated, may be driving the epithelial-to-mesenchymal transition (EMT) in a variety of cancers such as breast cancer (Mekhdjian *et al*., 2017).

Based on these previous results, we tested the hypothesis that a key mechanistic difference between GBM molecular subtypes is that proneural cells are slow migrating and mesenchymal cells are fast migrating. To address this question, we generated animal models recapitulating the transcriptomic signatures of human mesenchymal and proneural GBM in an immune competent background using perturbations of known GBM oncogenic pathways. Specifically, mesenchymal and proneural-like tumors were driven by SV40-large T (LgT) antigen, to mimic common inhibition of p53 and Rb signaling found in GBM (Ahuja, Sáenz-Robles and Pipas, 2005; McLendon *et al*., 2008), in combination with either NRAS^G12V^ (NRAS) or PDGFB (PDGF), respectively, which resulted in mesenchymal or proneural transcriptomic features with only a single genetic change required to switch subtypes in a wild type mouse background. As predicted, *CD44* expression was higher in NRAS-driven tumors and, consistent with our simulation predictions, *ex vivo* brain slice live imaging showed NRAS tumor cells migrate faster than PDGF tumor cells, and exhibit greater spreading, polarization, and force generation as well. Despite increased migration, the NRAS cohort had better survival than PDGF which was attributed to enhanced antitumoral immune response in NRAS tumors, consistent with increased immune cell infiltration in human mesenchymal GBM (Doucette *et al*., 2013; Wang *et al*., 2017, 2021; Hara *et al*., 2021). Consistently, NRAS-driven tumors had shorter survival in immunodeficient (Rag1^−^/^−^) mice compared to immunocompetent mice, supporting a role for the immune system in limiting tumor progression in mesenchymal tumors. Overall our work identified a clinically actionable difference in migration mechanics between GBM subtypes and establishes an integrated biophysical modeling and experimental approach to mechanically parameterize and simulate distinct molecular subtypes in preclinical models of cancer.

## MATERIALS AND METHODS

### Generation of mouse tumor models

All animal studies were conducted according to guidelines approved by the Institutional Animal Care and Use Committee at the University of Minnesota. All animals were housed in a daily monitored animal facility. Both immunocompetent FVB/NJ WT and immunodeficient FVB/N mice homozygous for the Rag1^tm1/Mom^ mutation (Rag1^−/-^) strains of mice were used in this study. Malignant gliomas were induced in neonatal mice by DNA plasmid injection into the right lateral ventricle as described previously (Wiesner *et al*., 2009; Calinescu *et al*., 2015). Briefly, neonatal mice were injected with 1 µg of plasmid DNA mixed with polyethyleneimine (jetPEI, Polyplus, Berkeley, CA), and 5% dextrose in a total volume of 2 µL at a rate of 0.7 µL/min. The following four plasmids were used (1:1:1:1) ratio: empty vector, pT2/C-Luc/PGK-SB100, pT/CMV-LgTAg-IRES-GFP, pT2/Cag-NrasV12 or pT2/Cag-mPDGF. Tumor initiation and detection require stable integration of all plasmids, ensuring that only cells with successful delivery and expression of full construct set give rise to detectable tumors. Animals were monitored daily for morbidity by bioluminescent imaging.

### Immunohistochemistry of mouse tumor sections

Formalin fixed and paraffin embedded (FFPE) mouse brain tissues were used to prepare 4 µm thick slides. FFPE tissue slides were stained with hematoxylin and eosin (H&E) or IHC using standard methods. Table S7 contains a list of antibodies and reagents used for antigen retrieval, blocking and detection.

### Quantification of IHC staining of mouse tumor sections

Immunohistochemistry data was quantified by counting the number of pixels in an image that were positively DAB stained. To avoid user bias and subjective counting, k-means clustering was used to identify pixels representing areas of positive DAB, hematoxylin staining, and background. In this implementation, every pixel in an analyzed image is assigned to one of four clusters, each cluster representing a different component in the image: positive DAB-stained areas (brown), positive hematoxylin counter-stained areas (blue), unstained tissue (light blue), and background glass slide (beige). Digital images of equal sizes (2000×2000 pixels) of DAB stained and hematoxylin counterstained tumor samples were converted from RGB to the HSV color model. Three fields were sampled from representative areas from each tumor.

Using a custom written MATLAB algorithm, user input is used to define areas representing the four components (positive DAB stain, hematoxylin counterstain, unstained tissue, and background glass slide). These points are used as the initial estimates for the centroid locations of each of the four clusters. The squared Euclidean distance between each pixel’s HSV coordinates and the HSV coordinates of each cluster’s centroid is computed. Each pixel is then assigned to the cluster with the minimum squared Euclidean distance to the cluster centroid. Cluster centroids are recalculated as the mean of the HSV coordinated of all current members. This process is iterated until the centroid of each cluster is stable. The number of pixels in the positive DAB-stained cluster is used to quantify percent positive pixels in Figure 7E-H.

### Transcriptional profiling of mouse tumors

Mice were euthanized in a CO_2_ chamber and perfused transcardially with isotonic saline. Mouse brains were extracted and GFP goggle (#FHS/EF-2G2; BLS-ltd, Budapest, Hungary) was used to dissect GFP-positive tumor tissues from NRAS (N=3) and PDGF (N=3) mouse brains. Two matched normal brain tissues were collected from brain regions away from the tumor and an additional normal brain tissue sample was collected from a health FVB adult mouse (N=3; Normal Brain Tissue; NBT). All samples were immediately placed in *RNALater* solution (Sigma, St. Louis, MO) for 24 hours then flash frozen in liquid nitrogen and stored in -80 °C for downstream processing. RNA extraction and sequencing were performed at the University of Minnesota Genomics Center (UMGC, Minneapolis, MN). RNA was extracted using *RNAeasy* Plus Universal Mini kit (Qiagen, Venlo, Netherlands) and libraries were prepared using *TruSeq* stranded mRNA (Illumina, San Diego, CA).

Next-generation sequencing was performed on the prepared RNA libraries using an Illumina HiSeq 2500 device in high output mode and generated 51 bp reads with an approximate depth of 20 million paired reads per sample. Mapping and expression calculations were generated using the rnaseq-pipeline of Gopher-pipelines (https://bitbucket.org/jgarbe/gopher-pipelines), which executed TopHat2 (Kim *et al*., 2013) and Cuffnorm (Trapnell *et al*., 2010) using the UCSC mm10 version of the mouse reference genome. Fastq files and the Cuffnorm output were deposited at Gene Expression Omnibus (GSE161154).

### Human GBM transcriptomic data

UNC RNASeqV2 level 3 expression (normalized RSEM) profiles of 171 samples (TCGA-GBM) were retrieved from Broad GDAC Firebrowse (Brennan *et al*., 2013). IDH status and subtype information were added to each sample based on the *Wang et al.* classification (Wang *et al*., 2017). For downstream analysis, 147 IDH-WT samples were used (57 Classical; 52 Mesenchymal; 38 Proneural).

### Clustering analysis of mouse and human expression profiles

To analyze the transcriptional profiles of mouse and human datasets, a value of 0.1 was added to all FPKM and RSEM values to minimize the impact of inaccurate low values (Scott *et al*., 2018). The expression data were log-transformed and mean-centered. Transcripts with a standard deviation greater than 1 were clustered using average linkage hierarchical clustering in MATLAB, with Pearson correlation used as the similarity metric. A custom MATLAB script was used to systematically identify transcriptional clusters within each dataset. For the mouse dataset, clusters were defined by a correlation threshold > 0.5 and a minimum of 100 transcripts. For the human dataset, a correlation threshold 0.2 and >100 transcripts were used. Fisher’s exact test was applied to compare cluster memberships in Figure 1C. All genes identified within each cluster are listed in Table S2.

**Figure 1.**
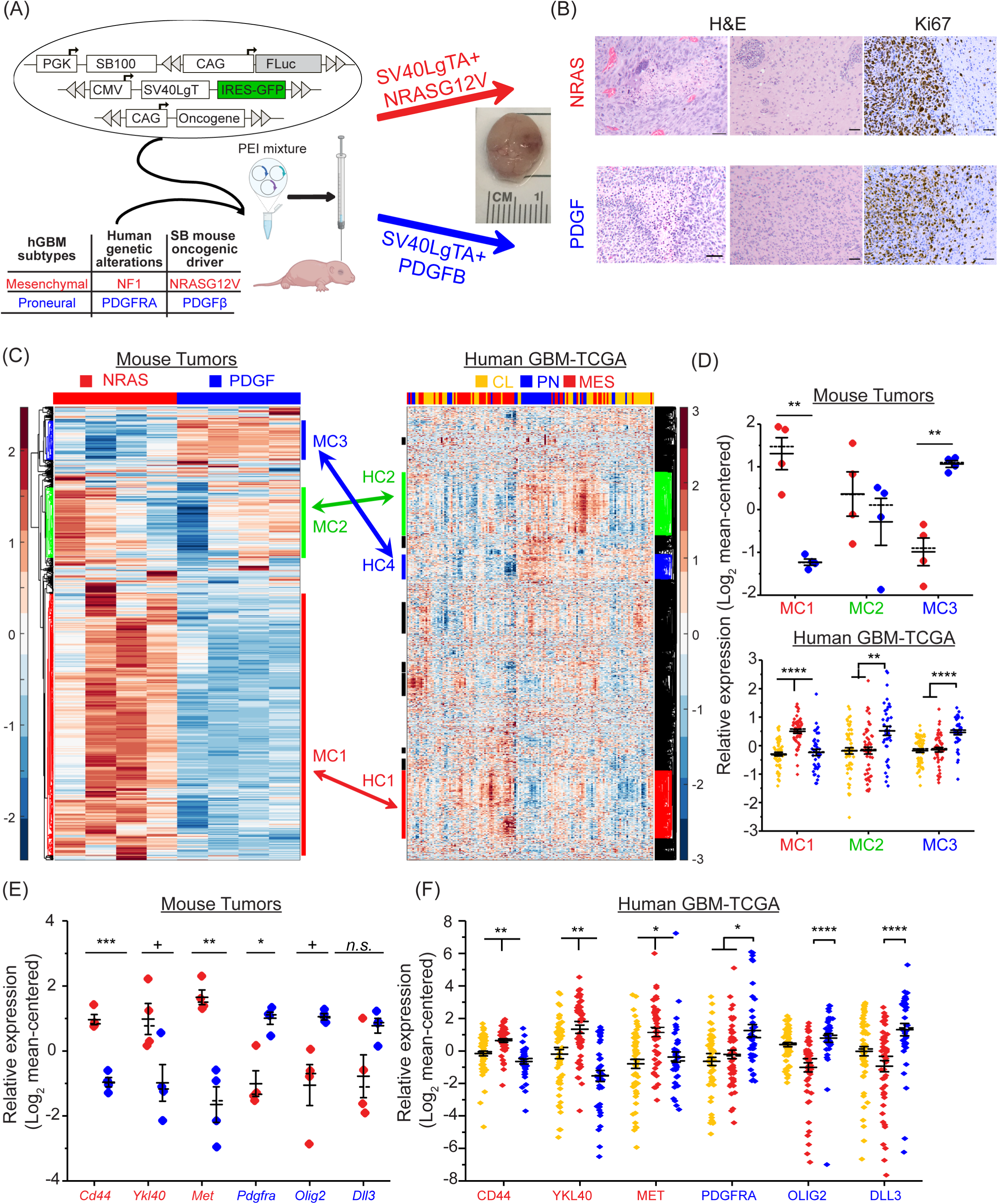
*De novo* induced GBM mouse models using immune competent mice recapitulate mesenchymal and proneural subtypes of human GBM. A) Schematic of mouse models. Plasmids encoding oncogenic drivers NRAS^G12V^ or PDGFβ in combination with SV40LgTA were injected into P1 FVB mice to induce mesenchymal and proneural high grade gliomas, respectively, Schematic was created with BioRender. B) H&E and Ki67 IHC staining of NRAS and PDGF tumor sections. Scale bar: 50 µm. C) Unsupervised hierarchical clustering of mRNA expression in induced mouse tumors and human IDH-WT GBM-TCGA. Arrows indicate conserved genes present in both mouse and human gene clusters as defined by systematic comparison of gene cluster membership between datasets. D) Quantification of relative expression of mouse gene clusters within the mouse dataset (top panel) and human GBM molecular subtypes (lower panel). E,F) Relative expression of key mesenchymal and proneural genes within mouse tumors (E) and human tumors (F). Solid and dashed lines represent mean and median values, respectively. Error bar represents S.E.M. + p <0.05, * p <0.01, ** p<0.001, *** p<0.0001, **** p<0.00001, Statistical significance was assessed using one-way ANOVA

**Figure 2.**
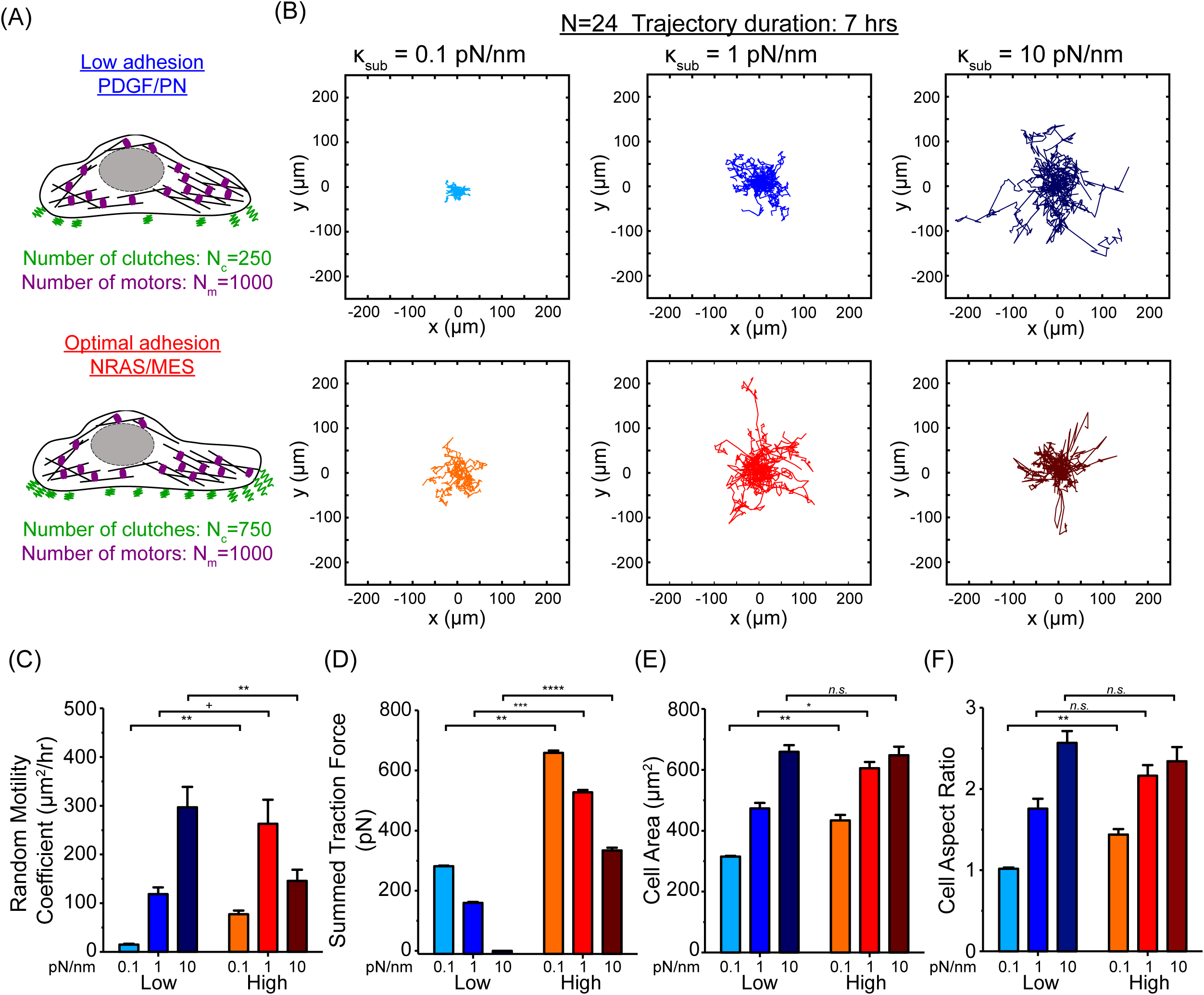
Simulations of cell migration as a function of subtype based on CD44-mediated cellular adhesive clutches. A) Schematic of proneural (low CD44) and mesenchymal (medium CD44, i.e. optimal) subtypes. B) Wind-rose plots of simulated cell trajectories on different stiffnesses (κ_sub_= 0.1, 1, 10 pN/nm). C) Simulated cell migration random motility coefficients show higher migration with increased (optimal) adhesion across a range of substrate stiffnesses. D) Cell summed traction force exhibits higher traction forces with increased (optimal) adhesion across all stiffnesses. E, F) Cell spread area and aspect ratio increase with increased adhesion on substrate stiffnesses of 0.1 and 1 pN/nm. Error bars are S.E.M. + p <0.05, * p <0.01, ** p<0.001, *** p<0.0001, **** p<0.00001, Statistical significance was assessed using the Kruskal–Wallis rank test with Dunn–Sidak correction

To quantify the relative expression of gene clusters (Figure 1D) and subtype-specific gene signatures (Figures S2C and S2D), we calculated the average expression of each gene set per sample. This was done by taking the simple average of mean-centered, log₂-transformed RNA expression values across all genes in the set. These per-sample averages were then plotted and used to calculate the mean relative expression for each mouse cohort and human GBM molecular subtype. Subtype gene signatures used for this analysis are listed in Table S3 (Wang *et al*., 2017). Table S3 also includes the subset of genes used in Figure S3, which were selected from the subtype gene signatures list based on our clustering criteria (standard deviation >1).

### Generation of mouse primary tumor lines

For each line, a tumor bearing mouse was euthanized and transcardially perfused with isotonic saline. Tumor tissue was collected and minced with a scalpel. Minced tumor fragments were incubated with PBS for 15 min at 37 °C. Tumor fragments were further dissociated by mixing them up and down using a 1000 μl micropipette. Finally, tumor suspension was passed through a 40 μm sterile cell strainer (Thermo Fischer Scientific, Waltham, MA) and filtrate was spun down and plated on a Matrigel (354230; Corning, Corning, NY) coated T-75 flasks (Corning, Corning, NY) using NSC media which consisted of DMEM/F12 (Gibco 11320033; Thermo Fischer Scientific, Waltham, MA) with 1X B-27 supplement (Gibco 12587010, Thermo Fischer Scientific, Waltham, MA) and 1X penicillin/streptomycin (Corning, Corning, NY). Twenty ng/ml EGF (PeproTech, Rocky Hill, NJ) and FGF (PeproTech, Rocky Hill, NJ) were added to the cell culture media every 2-3 days. Cells were cultured in a 37 °C 5% CO_2_ incubator. Once a confluent layer was achieved, cells were detached using 0.25% Trypsin EDTA (Corning, Corning, NY) and frozen down for later use.

Once tumor lines were established, cells were grown as neurospheres using NSC media and Ultra-Low Attachment 6-well plates (Corning, Corning, NY). Neurospheres were dissociated using accutase (Innovative Cell Technologies, San Diego, CA). In total, six different mouse primary tumor lines were established: three NRAS and three PDGF.

### Patient-derived xenograft (PDX) cell line culture

The patient-derived xenograft (PDX) cell lines were taken from the Mayo Clinic GBM PDX collection (managed by Dr. Jann Sarkaria, Mayo Clinic, Rochester, MN). Three mesenchymal PDX lines (GBM 16, 39 and 44) and three proneural PDX lines (GBM 64, 80 and 85) were selected to study their migration in organotypic mouse brain slice. Cells were cultured on Matrigel (354230; Corning, Corning, NY) coated tissue culture flasks in a 37 °C 5% CO_2_ incubator. NSC media was used to culture PDX cell lines and 20 ng/ml EGF and FGF were added to the cell culture media every 2-3 days.

### *Ex vivo* confocal imaging of tumor-bearing brain slices

Tumor-bearing mice were sacrificed when bioluminescence signals were around 5 × 10^7^ radiance (p/sec/cm^2^/sr). Mice were euthanized in a CO_2_ chamber and perfused transcardially with isotonic saline. Mouse brains were extracted and kept in chilled artificial cerebrospinal fluid (124 mM NaCl, 2.5 mM KCl, 2.0 mM MgSO_4_,1.25 mM KH_2_PO_4_, 26 mM NaHCO_3_, 10 mM glucose). Coronal brain slices of thickness 300 µm were prepared using a vibratome (Leica Biosystems, Buffalo Grove, IL). Only one slice was used for live-cell imaging. Isolectin GS-IB4 (Alexa Fluor 568 Conjugate; Molecular Probes, Eugene, OR) was used to label the vasculature.

Before imaging, the slice was washed and transferred into a No. 0 glass bottom 35 mm culture dish (P35G-0-20-C; MatTek, Ashland, MA) containing serum-free DMEM (Gibco, Thermo Fischer Scientific, Waltham, MA). A tissue culture anchor (SHD 42-15; Warner Instruments, Hamden, CT) was placed on top of the slice to prevent movement during imaging. The slice was then imaged on a Zeiss LSM 7 Live swept-field laser confocal microscope (Zeiss, Oberkochen, Germany) at 15-minute intervals for up to 20 hours in humidified 5% CO_2_ air at 37 °C. Images were collected with a 20x objective lens (Plan-ApoChromat 20X, 0.8 NA, Zeiss, Oberkochen, Germany). The number of Z stacks of several regions of interest was adjusted to ensure that the data acquisition of one frame in the time series was completed under 15 minutes (10-20 planes with 10 µm z-step was typically used). Maximum intensity projections from multiple Z stacks were used to generate 2D images for quantitative morphological and trajectory analysis. Images were registered by an affine transformation using ImageJ StackReg plug-in (École Polytechnique Fédérale De Lausanne) to account for stage drift and tissue relaxation during time-lapse imaging.

### Live-cell imaging of tumor cells in organotypic brain-slice culture

Healthy mouse brain slices were prepared using the same method as tumor-bearing brain slices above. For experiments using GFP*-*positive mouse primary tumor lines, neurospheres were dissociated using accutase (Innovative Cell Technologies, San Diego, CA). The protocol of grafting cancer cells into the brain slice was described in details in our previous publication (Liu *et al*., 2019). Briefly, after creating a single cell suspension, 300,000 cells in 3 mL of media were plated onto the brain slice. The cells were co-cultured with the brain slice for 4 hours at 37 °C and 5% CO_2_ before imaging to promote cell infiltration into the brain slice. Phenol-free NSC media +2% FBS (Gibco, Thermo Fischer Scientific, Waltham, MA) was used. The slices (Isolectin GS-IB4 stained) were washed several times using cell culture media and transferred into a No. 0 glass bottom 6-well plate (P06G-0-20-F; MatTek, Ashland, MA). The slices were imaged on a confocal microscope with a 10X objective lens (Plan-ApoChromat 10X, 0.45 NA, Zeiss, Oberkochen, Germany). Similar imaging protocol as mentioned above was applied. Each primary mouse line was imaged in one slice and was repeated at least twice except for mGBM3102 where only one successful slice was imaged and analyzed.

For PDX cells, 500,000 – 800,000 cells were stained using DiO membrane dye (V22886; Thermo Fisher Scientific, Waltham, MA) for 5 minutes and then washed twice before plating onto the brain slice inside a 35mm tissue culture dish. The grafting of the cells to the brain slice was similar to the mouse primary cells described above. The slices were imaged on a confocal microscope using a 20X objective lens (Plan-ApoChromat 20X, 0.8 NA, Zeiss, Oberkochen, Germany). Similar imaging protocol was applied. Experiment repeated 2-3 times on new set of slices each time.

For both mouse primary tumor cells and PDX cells, we also acquired the maximum intensity projections from multiple Z stacks and performed image registration for further analysis for cell migration and morphology. Images were registered by an affine transformation using ImageJ StackReg plug-in (École Polytechnique Fédérale De Lausanne) to account for stage drift and tissue relaxation during time-lapse imaging.

### Single cell tracking and morphology analysis

Single cell migration was tracked as previously described using a custom-written image segmentation algorithm in MATLAB (Bangasser *et al*., 2017; Klank *et al*., 2017a). Using cell centroid coordinates, the mean squared displacement (MSD) of the cell trajectories over time was calculated using the time interval overlap method (Dickinson and Tranquillo, 1993). To quantify the dispersion of cells, the MSD over time was used to calculate the random motility coefficient µ according to the equation (MSD(t)=4µt; assuming 2-D geometry). Using segmented cell regions, cell area and cell aspect ratio, defined as the ratio between the major and minor axis length of a fitted ellipse, were measured for each individual tracked cell. Average mean square displacement showed linear relation with time and random motility coefficients were calculated from each individual trajectory. Distributions of random motility coefficients, cell area and cell aspect ratio for the different conditions were compared using the Kruskal-Wallis test, which is a non-parametric rank-based test.

### Bioluminescence imaging and analysis

Animals were monitored for tumor development and progression using noninvasive bioluminescence imaging. Oncogene-injected animals were injected intraperitoneally with 100 µl of 28.5 mg/ml luciferin (GoldBio, St. Louis, MO) prior to imaging. Mice were then anesthetized using 3% isofluorane and imaged on an IVIS50 or IVIS100 instrument (Xenogen, Alameda, CA). Images were acquired ten minutes after injection with five minutes exposure time (Xenogen LivingImage Software, Alameda, CA). To avoid saturation, exposure time was reduced appropriately in fully grown tumors and accounted for in the analysis. BL images were processed using a custom written MATLAB algorithm where background signal was subtracted and pixels away from the tumor were set to zero. BL signal from each animal was then normalized to the initial time point when tumor was first detected.

### Quantification of proliferation of mouse primary tumor line

To measure proliferation rate, 200,000 cells from each line were plated into an ultra-low adhesion 6-well plate and grown as neurospheres. Growth factors were added every 2-3 days. At day six, neurospheres were dissociated and cells counted. After counting, the remaining cells were replated and resumed growing as neurospheres. Cells were also counted and replated at day nine and day 13. Experiment was repeated three times using each of five mouse primary tumor lines (three NRAS and two PDGF tumor lines). For each replicate, the average cell count for each cohort was calculated using the cell count from the different corresponding tumor lines.

### Traction force measurements

Traction force measurements of mouse primary tumor lines were performed using traction force microscopy on polyacrylamide gels embedded with 0.2 µm crimson fluorescent beads (Thermo Fischer Scientific, Waltham, MA) and coated with Type-I Collagen (Corning, Corning, NY). Collagen coated polyacrylamide gels of varying Young’s modulus were prepared as previously described (Wang and Pelham, 1998; Bangasser *et al*., 2017). Briefly, 0.7, 4.6, and 9.3 kPa polyacrylamide polymer mixture with fluorescent beads were cast onto a No. 0 glass bottom dish then coated with Type-I Collagen using Sulfo-SANPAH (Thermo Fischer Scientific, Waltham, MA). Mouse primary tumor cells were dissociated from neurospheres and plated on prepared gels at low density (1-5 cells/mm^2^) using NSC media +2% FBS.

To measure force transmission, Traction Force Microscopy (TFM) was performed as previously described (Bangasser *et al*., 2017). Briefly, Nikon TiE and Ti2 epifluorescence microscopes were used to image fluorescent bead positions before and after cell detachment via trypsin. A Zyla 5.5 sCMOS camera (Andor Technology, Belfast, United Kingdom) and a 40x/0.95NA Ph2 lens with 1.5x intermediate zoom (60x total magnification, 110 nm spatial sampling) was used. Cells were maintained at 37 °C and 5% CO_2_ for the duration of imaging using an Oko lab Bold Line top stage humidified incubator (Okolab, Ottaviano, Italy). At each stage position, a phase contrast image of the cell was acquired. Next, an image of fluorescent beads at the top surface of the gel was captured using a 575/25 nm LED and eGFP/mCherry filter set with LED fluorescence illumination from a SpectraX Light Engine (Lumencor, Beaverton, OR). Media in dishes was carefully removed, cells were detached with 0.25% trypsin/EDTA (Corning, Corning, NY), and fluorescence images of beads in the absence of cells were acquired at saved stage positions.

Using a previously described method (Bangasser *et al*., 2017), the displacement field was determined using particle image velocimetry (PIV) using the before and after bead images. A window size of 80-pixels (8.8 µm) square was used in PIV and a final lattice spacing of 20 pixels (2.2 µm) was achieved. Stress and displacement vectors were obtained by solving the inverse Boussinesq problem in Fourier space (Butler *et al*., 2002). By integrating the product of the stress and displacement vectors over the entire image, substrate strain energy was determined as previously described (Bangasser *et al*., 2017).

### Vascular deformation analysis

To visualize vessel deformation during tumor cell migration, a montage of representative time frames was generated from the vessel channel. In addition, a color-coded maximum-intensity projection over time was created, where the color of each pixel corresponds to the time point at which the maximum intensity occurred. In these projections, regions of the vessel that remained in the same position over the imaging period (i.e. were stationary) appeared gray, while regions that shifted position over time appear colored. Deformation was therefore defined as time-dependent displacement or bending of the vessel structure adjacent to the migrating cell. No additional quantitative metric was applied; the images are presented as a qualitative representation of vessel deformation (Fig. 5D, 5E; Video S3).

### Stochastic cell migration simulator

The previously described (Bangasser *et al*., 2017; Klank *et al*., 2017a) cell migration simulator (CMS v1.0) was used to simulate cells migration dynamics in response to changes in cell adhesion. The parameters used in the simulations are presented in Table S4. The number of adhesive clutches (*N_c_*) was adjusted to model the change in adhesion observed between PDGF/Proneural and NRAS/Mesenchymal tumors. N_c_ of 2500 and 7500 clutches were used to simulate PDGF/Proneural and NRAS/Mesenchymal tumor cells, respectively. Four hours of cell dynamics were simulated and the first hour was excluded from analysis to allow the system to reach steady state. Analysis was performed using a ten-minute sampling interval as previously described (Bangasser *et al*., 2017).

### Brownian dynamics tumor simulator (BDTS)

The Brownian dynamics tumor simulator was used as previously described with modifications (Klank, Rosenfeld and Odde, 2018; Ray *et al*., 2018). In the present study, we extended the BTDS to 3-dimentional tumors and incorporated immune cells’ dynamics as shown schematically in Figure 8A. Briefly, simulations started with 27 cancer cells, modeled as rigid sphere with radius (r_cancer_), placed in a 3×3x3 grid where the distance between each cancer cell (center-center) is 3*r_cancer_. In simulation including immune response, eight T cells, also modeled as rigid sphere with radius (r_CTL_), were included, and each was placed 1.5*r_cancer_ away from a randomly selected cancer cell. At each simulation time step of 1 min, cancer and T cells are allowed to move randomly and grow as spheres with a linear volumetric growth rate. Movement and growth are rejected if the newly assigned space is already occupied by a like cell (i.e. no-overlap enforced between cancer cells and between T cells). However, a cancer cell and T cell contact occurs when a proposed cell movement put the distance between cell centers less than or equal (r_CTL_+r_cancer_). The duration of the contact is (1/k_dissoc_), in this case 10 minutes. For every contact, both cancer cell and T cell take a “hit” that reduces their hit points (HP) by one. Both cancer cell and T cell have limited HP and once HP is depleted (equals 0), the cell dies or become exhausted. For NRAS simulations, T cells were added to the tumor simulator and T cell-mediated killing was simulated. For PDGF simulation, only cancer cells were simulated. Cancer cell motility was estimated from *ex vivo* brain slice imaging of tumor cells (Figure 3C) and proliferation rate was estimated from the *in vitro* proliferation of mouse primary tumor lines (Figure 7D). The rest of parameters were estimated based on previous published work or used as an adjustable parameter (see Table S6).

**Figure 3.**
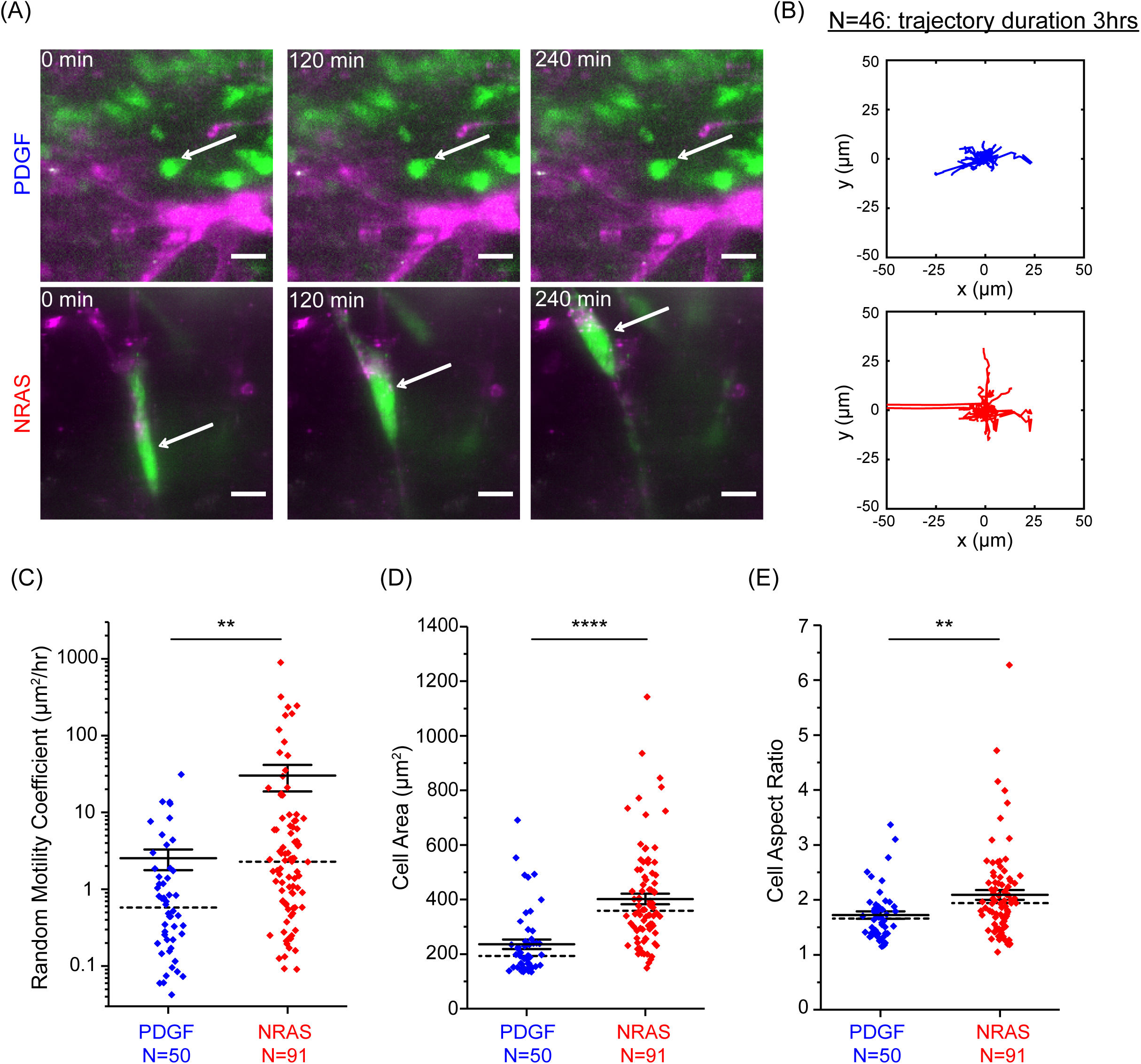
NRAS/Mesenchymal cells migrate faster, are more spread, and are more polarized than PDGF/Proneural cells in *ex vivo* tumor-bearing brain tissue. A) Representative *ex vivo* fluorescent montage of GFP-tagged tumor cells (green) and blood vessels staining using isolectin B4 (magenta). B) Wind rose plots of NRAS/Mesenchymal and PDGF/Proneural migrating tumor cell migration trajectories (N=46). C) NRAS/Mesenchymal cancer cell motility is faster than for PDGF/proneural cancer cells. D&E) NRAS/Mesenchymal cancer cells are more spread and polarized than PDGF/Proneural cells as evidenced by larger spread area and aspect ratio. Solid and dashed lines represent mean and median, respectively. Error bars are S.E.M. +p <0.05, * p <0.01, ** p<0.001, *** p<0.0001, **** p<0.00001. Statistical significance was assessed using Rank test, Kruskal-Wallis

**Figure 4.**
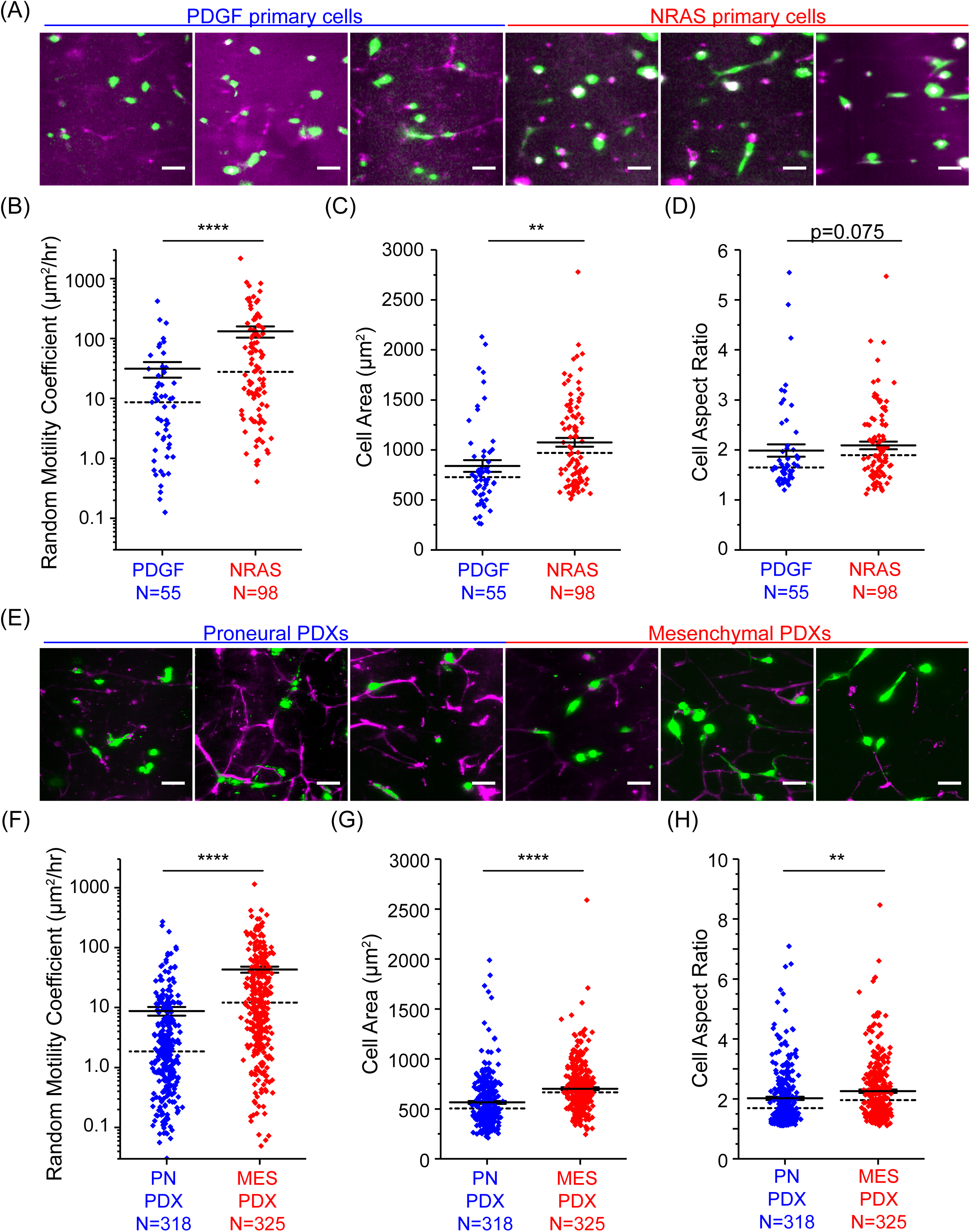
Cancer cell migration is subtype specific and independent of the tumor microenvironment and species. A) Six representative images of primary isolated cells from each of the two mouse tumor subtypes (3 NRAS and 3 PDGF). GFP-tagged tumor cells (green) and blood vessels staining using isolectin B4 (magenta). B) NRAS/Mesenchymal primary isolated tumor cancer cells migrate faster than PDGF/Proneural primary isolated cancer cells in normal mouse brain tissue as measured by random motility coefficient, see Video S2. C&D) Quantification of cell spread area and cell aspect ratio showing that NRAS/Mesenchymal cancer cells are more spread (C) and tend to be somewhat more polarized (D), p=0.075, than PDGF/Proneural cancer cells in healthy brain slices. E) Six representative images of proneural and mesenchymal PDX lines cells in healthy mouse brain tissue. Labeled tumor cells using DiO membrane dye (green) and blood vessel staining using isolectin B4 (magenta). F) Mesenchymal PDX cells migrate faster than proneural PDX cells in normal mouse brain tissue as measured by random motility coefficient, see Video S3. G&H) Quantification of cell spread area and cell aspect ratio showing that mesenchymal PDX cells are more spread (G) and are more polarized (H) than proneural PDX cells in healthy brain slices. Solid and dashed lines represent mean and median values respectively. Error bars are S.E.M. +p <0.05, * p <0.01, ** p<0.001, *** p<0.0001, **** p<0.00001. Statistical significance was assessed using Rank test, Kruskal-Wallis

**Figure 5.**
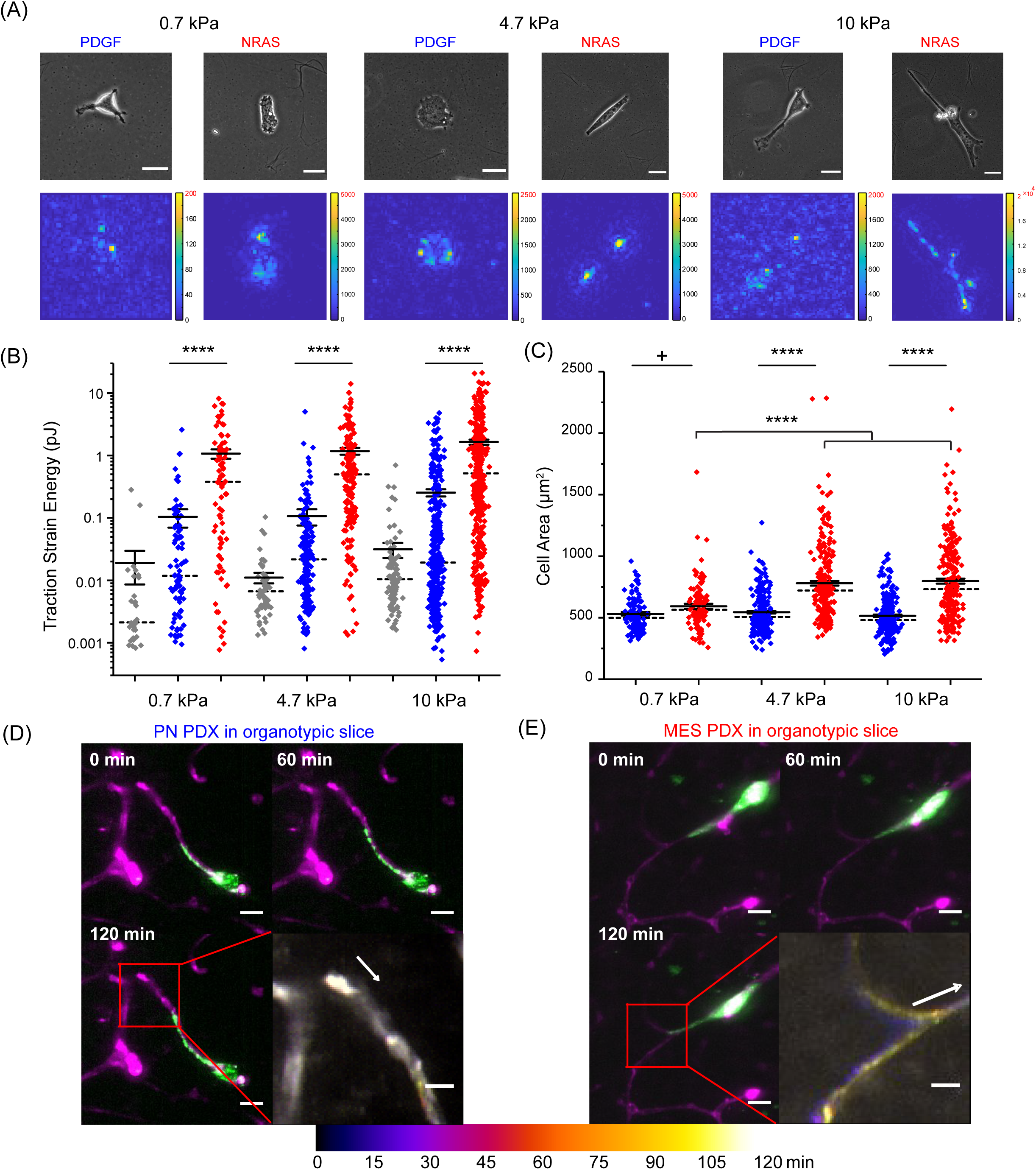
NRAS/Mesenchymal cancer cells generate larger traction forces *in vitro* on 2D hydrogels and *ex vivo* in brain slices than PDGF/Proneural cancer cells. A) Representative phase contrast images and traction force magnitude heatmaps on type I collagen-coated polyacrylamide hydrogels across different substrate stiffnesses. B) NRAS/Mesenchymal cancer cells (red) generate higher traction forces as measured by total strain energy relative to PDGF/Proneural (blue) cancer cells. Measurement noise is presented (grey) across all stiffnesses. C) Similar to mouse brain slices, NRAS/Mesenchymal cancer cells (red) cultured on collagen-coated polyacrylamide hydrogels have larger spread area than PDGF cancer cells (blue). D &E) Tissue deformation visualized using temporal-color coding using 2 hours time-lapse and 15 minutes time interval. (D–E) Representative montage of three time points and corresponding color-coded maximum projection over time. Color indicates the time point at which the maximum intensity occurred; gray indicates no displacement (D) Proneural PDX cells and (E) Mesenchymal PDX cells. generate larger deformations of mouse brain capillaries (E) than proneural PDX cells shown in (D). Arrows point to migrating cell direction. Scale bar = 20 µm, and inset scale bar = 10 µm. Solid and dashed lines represent mean and median values, respectively. Error bars are S.E.M. +p <0.05, * p <0.01, ** p<0.001, *** p<0.0001, **** p<0.00001. Statistical significance was assessed using Rank test, Kruskal-Wallis.

### Statistical analysis

Fisher’s exact test was used to compare the mouse and human transcriptomic clusters. One-way analysis of variance was used to compare transcript levels. Analysis of covariance (ANCOVA) was used to compare between the two regression lines in Figure 5C. Rank test, Kruskal-Wallis, one-way analysis of variance was used to compare single cell behaviors and IHC quantifications. Where appropriate, a subsequent Dunn-Sidak test for multiple comparisons was performed.

## DATA AND CODE AVAILABILITY

All data and codes are available on the Odde laboratory website (oddelab.umn.edu) or on request from the corresponding author. Fastq files and the Cuffnorm output were deposited at Gene Expression Omnibus (GSE161154).

## RESULTS

### Genetically induced high-grade glioma mouse models recapitulate the transcriptomic signatures of mesenchymal and proneural GBM

To characterize the mechanics of GBM subtypes, we utilized the *Sleeping Beauty* (SB) transposon-based gene transfer system to induce high grade gliomas in immunocompetent FVB/NJ-strain mice (Wiesner *et al*., 2009; Calinescu *et al*., 2015; Koschmann *et al*., 2016; Klank *et al*., 2017a; Núñez *et al*., 2019). Plasmid constructs carrying SB transposons with oncogenic driver transgenes (SV40-LgTA+NRAS^G12V^ or SV40-LgTA+PDGFB; here termed NRAS and PDGF, respectively) were used to model mesenchymal and proneural GBM tumors, respectively (Figure 1A). Additional plasmids encoding firefly luciferase and green fluorescent protein (GFP), as well as the SB transposase, were co-injected to enable stable genomic integration, confirm successful gene transfer, and allow monitoring of tumor development and growth via bioluminescence imaging (BLI) and single-cell tracking using fluorescence microscopy. Notably, tumor initiation and detection require stable integration of all plasmids, ensuring that only cells with successful delivery and expression of full construct set give rise to detectable tumors. Similar to human GBM, histological sections from these tumors exhibited highly mitotic tumor cells, necrosis, anaplasia, and perivascular infiltration and proliferation (Figure 1B).

To assess whether the NRAS and PDGF tumors recapitulated the mesenchymal and proneural subtypes, respectively, we performed cross-species transcriptomic analysis using bulk RNA sequencing data from mouse and human tumors. Bulk RNA sequencing was performed on tumor tissues from both cohorts and on normal brain tissues (NBT) (NRAS N=4, PDGF N=4, and NBT N=3). IDH-WT human GBM transcriptomic profiles were retrieved from Broad GDAC Firebrowse (Brennan *et al*., 2013). Unsupervised hierarchical clustering of the mouse dataset (Table S1) revealed clear differences between normal tissue and tumor tissue and between NRAS and PDGF tumors (Figure S1A). Not surprisingly, gene ontology enrichment analysis, using *EnrichR* (Kuleshov *et al*., 2016), showed an enrichment of cell cycle related processes in tumor tissue specific gene cluster (817 genes) (Figure S1B) and neuronal processes in normal tissue cluster (1722 genes) (Figure S1D). Interestingly, the NRAS-specific cluster (1327 genes) was enriched with cytokine-mediated signaling and inflammatory response processes (Figure S1C).

To determine whether any variations observed between the two mouse cohorts were also present in human tumors, unsupervised hierarchical clustering was performed on both mouse and human tumor datasets and clusters were independently identified in both datasets (Table S2). Three and 10 gene clusters were identified in both mouse and human datasets, respectively, as shown in Figure 1C. We identified mouse cluster MC1 (n=1534 genes) as being significantly enriched in genes found in human cluster HC1 (n=1186 genes, p < 1×10^−15^), MC2 (n=414 genes) is significantly enriched in genes found in HC2 (n=1098 genes, p < 1×10^−15^), and MC3 (n=232 genes) is significantly enriched in genes enriched in HC4 (n=432, p < 1×10^−15^). These results show that conserved transcriptomic patterns distinguish subtypes of both mouse and human tumors.

To assess whether the transcriptional patterns present in the mouse tumor models represent previously described GBM subtypes, we compared the expression of identified mouse gene clusters within human GBM subtypes. We found that MC1, which is enriched in NRAS tumors, is significantly enriched in human mesenchymal GBM relative to proneural and classical GBMs (Figure 1D). In contrast, MC3, which is enriched in PDGF tumors, is significantly enriched in proneural GBM relative to mesenchymal and classical GBMs (Figure 1D). Furthermore, we found the expression of known mesenchymal and proneural genes and gene signatures are relatively elevated in NRAS and PDGF tumors, respectively (Figure 1E, 1F and Figure S2). These results demonstrate that NRAS and PDGF tumors transcriptionally resemble mesenchymal and proneural GBMs, respectively, consistent with established subtype markers and gene signatures defined (Phillips *et al*., 2006; Verhaak *et al*., 2010; Wang *et al*., 2017).

### Motor-clutch modeling of cell migration predicts NRAS/Mesenchymal tumor cells will migrate faster, have larger cell spread area, and generate more force than PDGF/Proneural tumor cells

To examine tumor cell migration, we used our cell migration simulator (Bangasser *et al*., 2017; Klank *et al*., 2017a) to predict migration phenotypes in response to gene expression changes. The cell migration simulator is based on the motor-clutch model which incorporates actin-based protrusion dynamics, mass conservation, and force balances to reproduce cell polarization and random motility in 1D and 2D compliant microenvironments (Chan and Odde, 2008; Bangasser, Rosenfeld and Odde, 2013; Bangasser *et al*., 2017; Klank *et al*., 2017a; Estabridis *et al*., 2018; Prahl *et al*., 2018, 2020; Hou *et al*., 2019; Liu *et al*., 2019). The number of adhesion/clutches and motors are key determinants of cell migration, with a relative balance being essential for efficient migration (DiMilla, Barbee and Lauffenburger, 1991; Bangasser, Rosenfeld and Odde, 2013; Bangasser *et al*., 2017). Using a representative set of 54 cell migration genes previously identified to be expressed in the human U251 GBM cell line (Bangasser *et al*., 2017), NRAS tumors showed significantly increased expression of adhesion and adapter genes (Figure S3A). A similar pattern was observed in MES relative to PN tumors in the TCGA-GBM dataset (Figure S3B). Both NRAS and MES tumors upregulated *CD44* and its cognate adhesion adapter gene moesin (*MSN*), which mechanically links the CD44 cytoplasmic tail to F-actin (Tsukita *et al*., 1994; Legg and Isacke, 1998; Yonemura, 1998; Ponta, Sherman and Herrlich, 2003; Toole, 2009; Fehon, McClatchey and Bretscher, 2010; Freeman *et al*., 2018). Notably, the levels of myosin motor genes were not differentially expressed in the mouse dataset, while, in the human dataset, *MYH9* and *MYO1C* were modestly upregulated in MES tumors but to a lesser degree than adhesion molecules (Figure S3B). These results suggest NRAS/Mesenchymal tumor cells have a higher number of adhesion/clutches than PDGF/Proneural tumor cells and little to no change in the number of motors.

Based on these results, we simulated the effect of *CD44* expression level on cell migration by adjusting the number of adhesion bonds (number of clutches, *N_c_*) in the model (Bangasser *et al*., 2017; Klank *et al*., 2017a). We used low clutches relative to motors (low adhesion) to simulate PDGF/Proneural cells and a medium level of clutches that balanced the number of motors (optimal adhesion) to simulate NRAS/Mesenchymal cells (Figure 2A).

Our simulations show that lowering the number of adhesions, representing the PDGF/Proneural case, results in reduced cell migration, force transmission, cell spread area, and cell polarization due to an insufficient number of clutches relative to the number of motors, as shown in Figure 2B-F. In contrast, when clutch and motor numbers are balanced, representing the NRAS/Mesenchymal case, simulated cells recover their ability to migrate, transmit forces, spread, and polarize across a range of substrate stiffnesses. Consequently, simulation results predict that NRAS/Mesenchymal tumor cells will migrate faster than PDGF/Proneural tumor cells due to increase of adhesion (i.e. CD44 expression) rather than small differences in motor gene expression (Figure S3A and S3B). In addition, with a higher number of clutches and a balanced motor-clutch ratio, NRAS/Mesenchymal tumor cells are predicted to generate higher force, spread more, and display greater polarization

### NRAS/Mesenchymal tumor cells migrate faster than PDGF/Proneural tumor cells in brain tissue

To test our model prediction that NRAS/Mesenchymal cells migrate faster than PDGF/Proneural cells, we performed live cell imaging on tumor bearing mouse brain slices using confocal microscopy. Time-lapse images of GFP-positive tumor cells were used to track single cell migration and generate single cell trajectories. As shown in Figure 3A, 3B and Video S1, NRAS/Mesenchymal tumor cells appeared qualitatively to move farther, have larger spread area, and polarize to a greater extent than PDGF/Proneural tumor cells. Quantitative analysis of single cell trajectories confirmed that NRAS/Mesenchymal tumor cells have a higher random motility coefficient than PDGF/Proneural tumor cells (30.1 µm^2^ hr^−1^ *vs* 2.5 µm^2^ hr^−1^, p<0.001; see Figure 3C and S4A). In addition, morphological analysis of tumor cells revealed cell spread area and cell aspect ratio (i.e. polarization) are also higher in NRAS/Mesenchymal tumor cells than PDGF/Proneural tumor cells (406.6 µm^2^ vs 235.8 µm^2^, p<0.00001 and 2.1 *vs* 1.7, p<0.001, respectively, see Figure 3D, 3E, S4B and S4C). As predicted by our modeling, and the hypothesis that NRAS/Mesenchymal has balanced motors and clutches while PDGF/Proneural lacks sufficient clutches, we find NRAS/Mesenchymal cells migrate faster, are more spread, and are more polarized than PDGF/Proneural cells.

### Migration phenotype is species and tumor microenvironment independent

To determine whether migration phenotype is cancer cell intrinsic as predicted by our modeling and not due to microenvironment differences, we generated three primary mouse lines grown as neurospheres from each cohort to investigate their migration phenotype outside their tumor microenvironment. Organotypic mouse brain slice culture was used to image tumor cell migration in healthy mouse brain slices (Liu *et al*., 2019). Dissociated mouse tumor cells were plated and allowed to invade and migrate in healthy mouse brain slices. Figure 4A shows representative fluorescence images of primary isolated cells in organotypic slice culture. Time-lapse imaging was used to track single cells and quantify their migration rates. Consistent with the *ex vivo* migration in intact tumor-bearing brain slices, random motility coefficient in normal mouse brain tissue is higher for primary NRAS/Mesenchymal tumor cells than primary PDGF/Proneural tumor cells (131.4 µm^2^ hr^−1^ *vs* 31.3 µm^2^ hr^−1^, p<0.00001; Figure 4B, S4D and Video S2). The area of cell spreading is also higher in primary NRAS/Mesenchymal tumor cells than PDGF/Proneural (1075.1 µm^2^ *vs* 838.8 µm^2^, p<0.001; Figure 4C and S4E). Furthermore, cell aspect ratio, the ratio of major and minor axis of a fitted ellipse, was trending higher in NRAS/Mesenchymal tumor cells than PDGF/Proneural but did not reach statistical significance (Figure 4D and S4F).

To assess the relevance of these results to human GBM, we tested the migration phenotype of six patient-derived xenograft (PDX) lines (three mesenchymal and three proneural) using the organotypic mouse brain slice culture (Table S5). Figure 4E shows representative fluorescence images of PDX cells in organotypic slice culture. We found mesenchymal PDX cells migrate faster than proneural PDX cells (43. µm^2^ hr^−1^ *vs* 8.8 µm^2^ hr^−1^, p<0.00001; Figure 4F, S4G and Video S3). In addition, similar to our mouse models, mesenchymal PDX cells have larger area of cellular spreading and aspect ratio relative to proneural PDX cells (701.7 µm^2^ vs 564.6 µm^2^, p<0.00001 and 2.3 *vs* 2.0, p<0.001, respectively, see Figure 4G, S4H, 4H and S4I).

### Traction strain energy is larger for NRAS/Mesenchymal cells than for PDGF/Proneural cells, consistent with model predictions

In addition, cell migration simulations predict NRAS/Mesenchymal tumor cells would have increased force generation as a result of higher number of clutches resulting in balanced myosin motors and clutches, relative to PDGF/Proneural tumor cells which would have insufficient clutches relative to motors (Figure 2D). Using the primary isolated mouse lines, traction force microscopy was used to measured traction strain energy generated by tumor cells on polyacrylamide hydrogels coated with type-I collagen (Butler *et al*., 2002; Bangasser *et al*., 2017). Consistent with model predictions, NRAS/Mesenchymal tumor cells generate higher traction strain energy than PDGF/Proneural tumor cells across different substrate Young’s moduli (Figure 5A and 5B). Cell spread area is also higher in NRAS/Mesenchymal than PDGF/Proneural on polyacrylamide hydrogels (Figure 5C). In addition, NRAS/Mesenchymal cells exhibit stiffness sensitive cell spreading; cells on stiff substrate (4.6 and 9.3 kPa) were more spread than on soft substrate (0.7kPa, p <0.00001; see Figure 5C). Furthermore, we also examined force generation of mesenchymal and proneural PDX cells in mouse brain slices. Qualitative analysis of vasculature deformation is consistent with the model prediction that mesenchymal PDX cells generate larger deformations relative to proneural PDX cells. These deformations are identified from max-intensity projections of the vessel channel over time, where stationary regions remain gray and regions that shift over time appear colored. In this representation, deformation corresponds to a time-dependent displacement or bending of the vessel near the migrating tumor cell as shown in Figure 5D, 5E and Video S3.

### NRAS/Mesenchymal mice have better survival and slower tumor growth rate

Since NRAS/Mesenchymal cells have nearly optimal *CD44* expression, and therefore higher motility compared to PDGF/Proneural cells which have a suboptimal low level of *CD44* expression (Klank *et al*., 2017a), we asked whether the differences in migration rate, morphology and force generation correlate with disease progression and survival. Specifically, based on the faster migration in the NRAS/Mesenchymal cohort, we expected that these mice would progress faster and die sooner than PDGF/Proneural mice. To test this hypothesis, we measured survival times of tumor bearing mice and found that, opposite to our expectation, the NRAS/Mesenchymal cohort had better median survival than PDGF/Proneural cohort (NRAS N=21, PDGF N=24, 65 days vs. 35 days, log-rank test, p<0.0001; Figure 6A). To explain the difference in survival, we quantified *in vivo* tumor growth using bioluminescence imaging (BLI) of tumor-bearing mice. Consistent with their shorter survival, we found PDGF tumors grew twice as fast as NRAS tumors (Slopes: 0.127±0.01191 vs. 0.0716±0.00343 p <0.001, Figure 6B and 6C). Using mouse tumor neurospheres, we quantified mouse primary tumor cell line proliferation rates *in vitro* and found no significant difference between NRAS/Mesenchymal and PDGF/Proneural cells (Figure 6D). These results imply that an additional factor, besides proliferation or migration, enables the NRAS/Mesenchymal mice to live longer and their tumors to grow slower *in vivo* than PDGF/Proneural mice.

**Figure 6.**
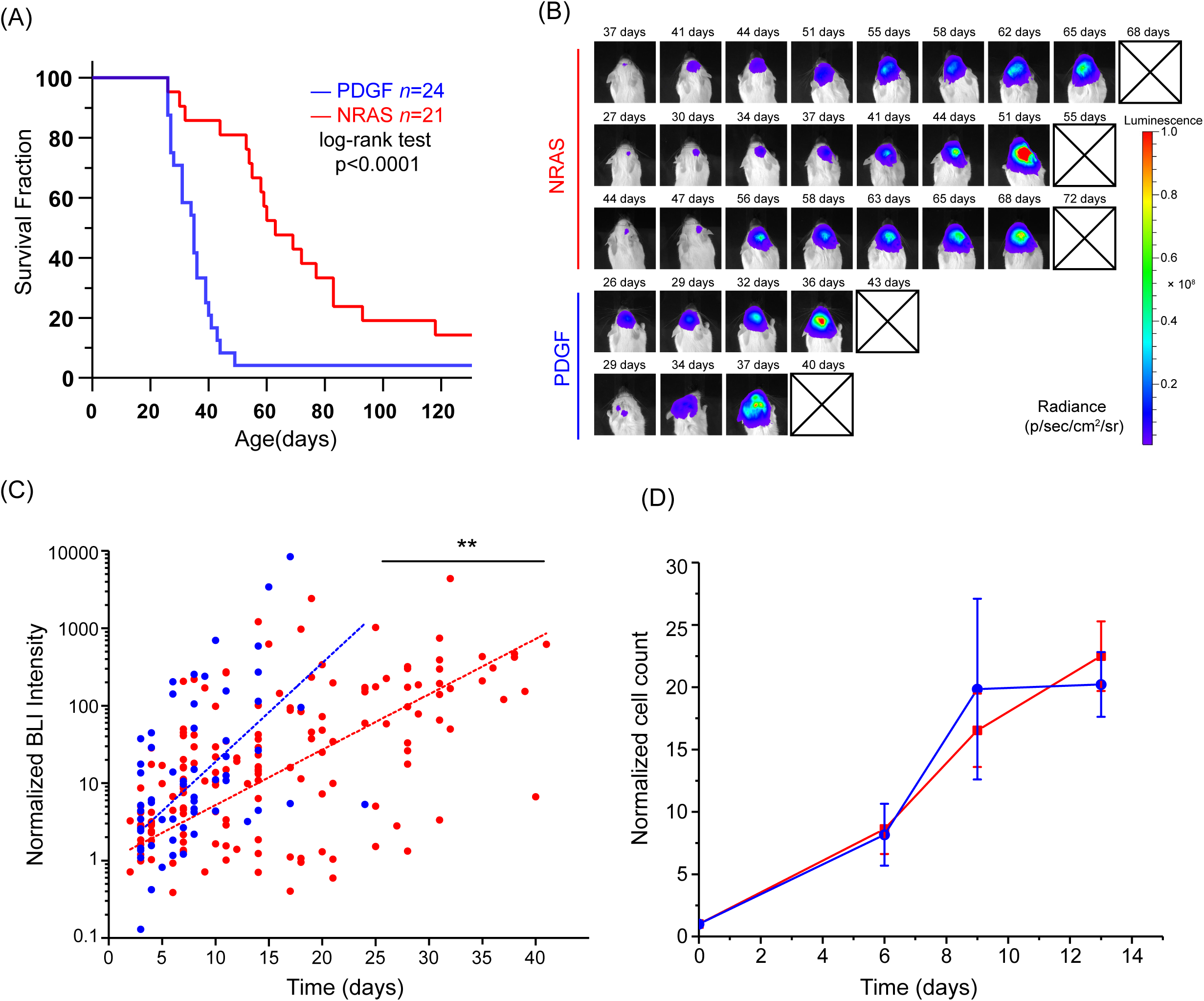
NRAS/Mesenchymal mice live longer than PDGF/Proneural mice. A) Kaplan-Meier plot of animal survival for NRAS/Mesenchymal and PDGF/Proneural tumor-bearing mice shows extended survival of the NRAS/Mesenchymal cohort. Log rank Mantel–Cox test p<0.00001. B) Bioluminescence imaging (BLI) of NRAS/Mesenchymal mice shows slower growing tumors relative to PDGF/Proneural mice. C) Quantification of BLI integrated intensity for the two cohorts. Normalized BLI intensity for all data points and linear fits for each cohort. PDGF/Proneural slope=0.127±0.01191 and NRAS/Mesenchymal slope=0.0716±0.00343, D) Ki67 IHC staining of mouse tumor sections, scale bar = 50 µm. E) Normalized cell count of primary isolated mouse tumor cells showing no significant difference in proliferation rate *in vitro* using neurosphere culture. Solid and dashed lines represent mean and median values, respectively. Error bars are S.E.M. +p <0.05, * p <0.01, ** p<0.001, *** p<0.0001, **** p<0.00001. Statistical significance was assessed using ANCOVA to compare two regression lines

**Figure 7.**
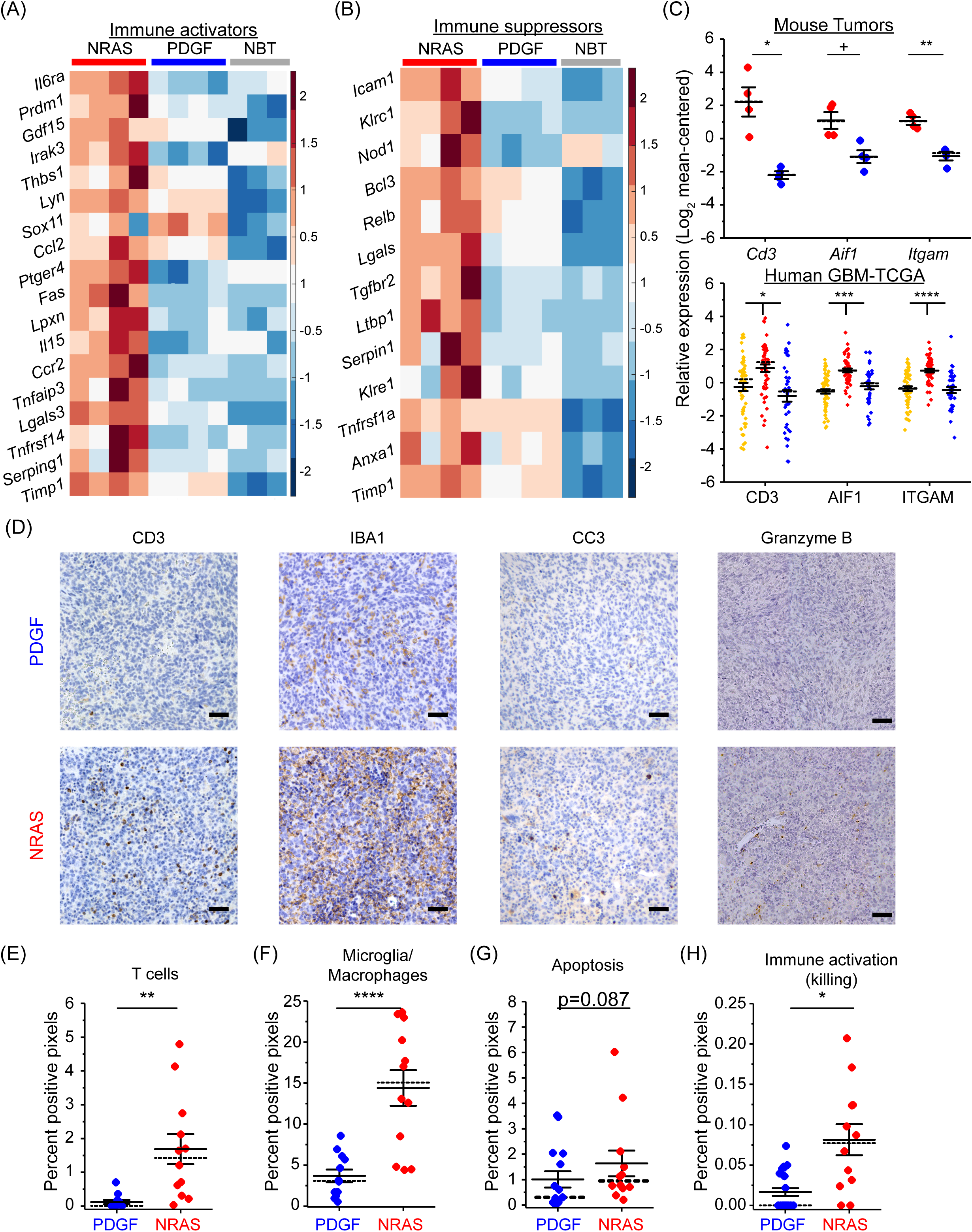
NRAS/Mesenchymal tumors are immunologically “hot” and PDGF/Proneural tumors are immunologically “cold,” consistent with human GBM subtypes. A, B) Clustering analysis of immune activators and suppressors previously reported in human GBM (Doucette *et al*., 2013). Similar to mesenchymal GBM, NRAS/Mesenchymal tumors have elevated immune activator and suppressor expression relative to normal brain tissue and PDGF/Proneural tumors. C) Expression of immune cell surface marker genes in mouse (top) and human (bottom) tumors shows elevated expression in NRAS/Mesenchymal tumors and human mesenchymal GBM tumors relative to PDGF/Proneural and human proneural GBM tumors. D) Immunohistochemistry (IHC) confirms elevated immune cell infiltration in NRAS tumors and associated elevation of immune-mediated killing of tumor cells. E,F,G,H) Quantification of IHC images using a k-means clustering algorithm. Solid and dashed lines represent mean and median values, respectively. Error bars are S.E.M. +p <0.05 * p <0.01, ** p<0.001, *** p<0.0001, **** p<0.00001. Statistical significance was assessed using one-way ANOVA

**Figure 8.**
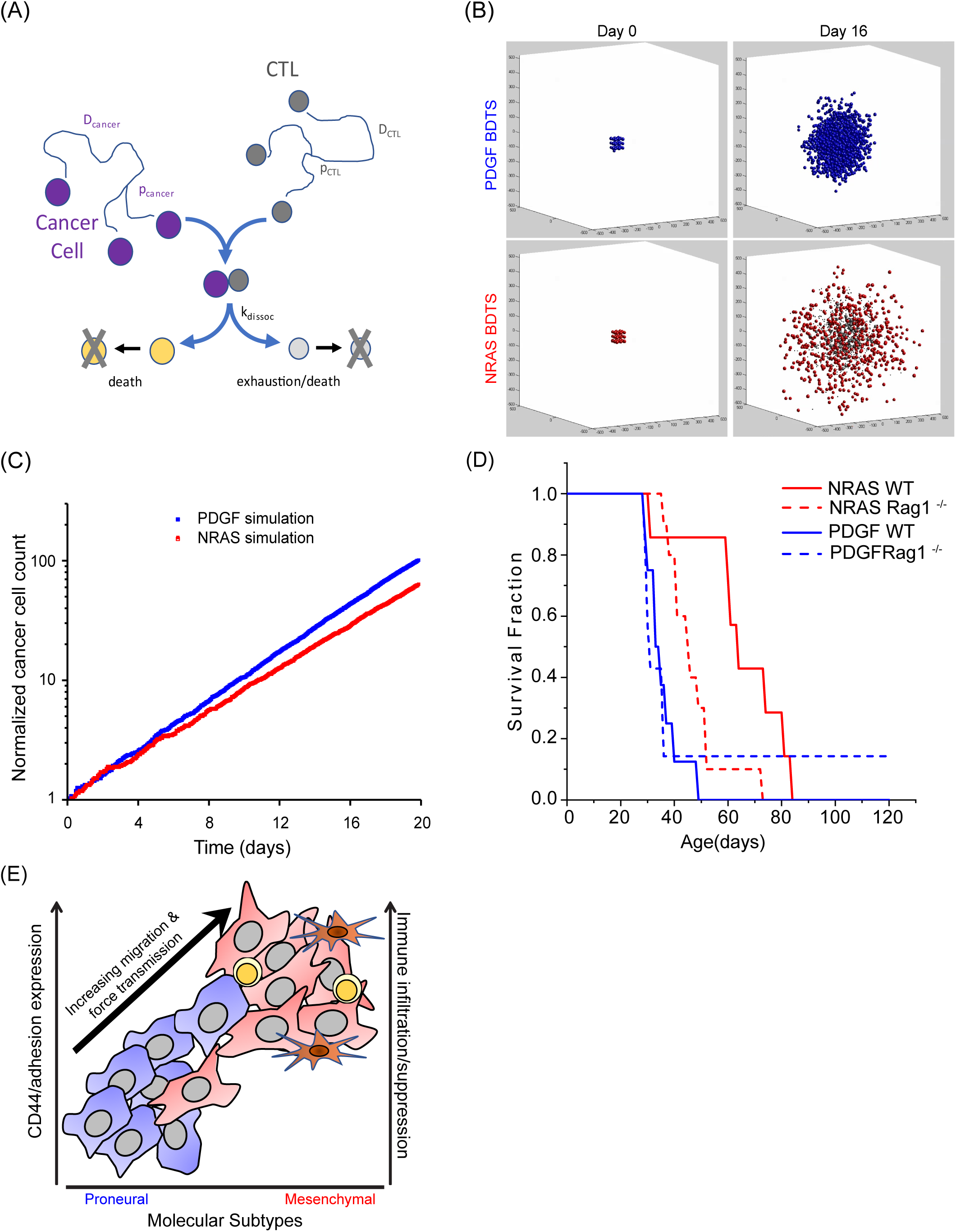
Brownian dynamics tumor simulator (BDTS) of 3D NRAS/Mesenchymal and PDGF/Proneural tumors. A) Schematic showing a diagram of the model. B) Simulator output at day 0 and day 16 for PDGF and NRAS simulations. In NRAS simulations, dead cancer cells are larger grey spheres and T cells are smaller black spheres, see Video S4. C) Cancer cell count over time showing the difference in tumor growth in the presence of T cells with simulated PDGF/Proneural tumors growing faster than NRAS/mesenchymal tumors. D) Kaplan-Meier plot for NRAS/Mesenchymal and PDGF/Proneural tumor-bearing mice in wild-type (WT) versus immunodeficient (Rag1^−/-^) background. NRAS/Mesenchymal tumors show significantly reduced survival in Rag1-/-mice compared with WT (Log rank Mantel–Cox test p=0.012). PDGF/Proneural survival is not affected. E) Cartoon summary of the main findings depicting the relationship between molecular subtypes, cellular adhesion, cell migration, and immune response.

### NRAS/Mesenchymal mice have increased immune response relative to PDGF/Proneural mice

Because mesenchymal GBMs are known to be immunologically “hot” – which presumably confers a survival benefit due to an antitumoral immune response- relative to immunologically “cold” proneural GBMs (Doucette *et al*., 2013; Wang *et al*., 2017; Neftel *et al*., 2019), we assessed the extent to which mouse NRAS/Mesenchymal tumors induce an immunological response relative to PDGF/Proneural tumors and normal brain tissues. Using our mouse transcriptomic dataset and previously published GBM immune gene sets (Doucette *et al*., 2013), NRAS/Mesenchymal tumors were found to have increased expression of both immune activators and suppressors gene signatures similar to human mesenchymal GBM (Doucette *et al*., 2013), as shown in Figure 7A and 7B. Specifically, expression of immune cell marker genes such as *Aif1*, *Itgam* (microglia/macrophages) and *Cd3* (T cells) are elevated in NRAS/Mesenchymal tumors relative to PDGF/Proneural tumors (Figure 7C upper panel) and in mesenchymal GBMs relative to proneural and classical GBMs (Figure 7C lower panel). Elevated expression of immune cell markers is indicative of increased immune cell infiltration in mesenchymal tumors. Consistent with the transcriptomic findings, IHC staining of NRAS/Mesenchymal and PDGF/Proneural tumor sections revealed significant levels of immune cell infiltration, including both microglia/macrophages and T-lymphocytes, in NRAS/Mesenchymal but much less so in PDGF/proneural tumors (Figure 7D). Furthermore, increased immune infiltration and activity in NRAS/Mesenchymal tumors was accompanied by increased cell killing as measured by granzyme B and cleaved caspase-3 staining (Figure 7D). Image clustering analysis was used to quantify CD3, IBA1, granzyme B, and cleaved caspase-3 staining, and statistically significant differences were observed between NRAS/Mesenchymal and PDGF/Proneural cohorts (Figure 7E-H). This anti-tumor response was also evident in the three instances where NRAS mice developed tumors and tumor regression was observed (Figure S5); whereas, in long-surviving PDGF cohort, there was no evidence of tumor regression. Despite the anti-tumor immune response observed in NRAS/Mesenchymal tumors, transcriptomic analysis also revealed elevated relative expression of immune checkpoint genes including PDL1, CTLA4, and CD200R1 (Figure S6). Thus, the NRAS/mesenchymal tumors, like human mesenchymal GBMs, are immunologically “hot” with evidence of both immune activation and immune suppression, as well as evidence of cell killing. Altogether, the enhanced immune cell-associated tumor cell killing provides a mechanism by which survival is extended in NRAS/Mesenchymal tumors despite their enhanced migration speeds relative to PDGF/Proneural tumors.

### Brownian dynamics simulations and immunodeficient tumors model explain NRAS/Mesenchymal and PDGF/Proneural tumor progression

To quantitatively describe the interplay between tumor cell migration and proliferation and anti-tumoral immune response in tumor growth, we developed a three-dimensional (3D) Brownian dynamics tumor simulator (BDTS) based on our original 1D Brownian dynamics simulator (Klank, Rosenfeld and Odde, 2018; Ray *et al*., 2018). The simulator takes into account anti-tumoral immune cells that infiltrate tissue, migrate, proliferate, encounter cancer cells, deliver cytotoxic agents, dissociate from cancer cells, undergo exhaustion, and, eventually, undergo apoptosis (or egress to lymphatics) as shown in Figure 8A. At the same time, cancer cells migrate, proliferate and undergo CTL-mediated death in the presence of anti-tumoral immune cells in the case NRAS/Mesenchymal tumors. In the case of PDGF/Proneural tumors, no anti-tumoral immune cells were simulated. Figure 8B shows simulation output at day 0 and day 16, which showed the observed behavior of overall faster growth of PDGF/Proneural tumors. In the NRAS/Mesenchymal tumor simulations, cancer cells appear more dispersed, whereas, in PDGF/Proneural simulations, cancer cells are less dispersed. Simulated tumor growths were plotted in Figure 8C and shown in Video S4. Using the parameters in Table S6, including the experimentally observed single cell migration speeds and neurosphere proliferation rates, simulated tumors qualitatively recapitulate the *in vivo* growth profile of NRAS/Mesenchymal and PDGF/Proneural tumors without parameter adjustment (Figure 8C).

Lastly, given the strong immune gene signature and immune cell infiltration observed in NRAS/Mesenchymal tumors, we sought to determine whether this immune response functionally impact tumor progression. To test this, NRAS and PDGF tumors were induced in an immunodeficient (Rag1^−^/^−^) mice to assess the role of the adaptive immune system in controlling NRAS/Mesenchymal vs. PDGF/Proneural tumor growth. Compared to immunocompetent wild-type (WT) mice, NRAS tumor-bearing Rag1^−^/^−^mice exhibited significantly shorter survival (WT *n*=7, Rag1^−^/^−^ *n*=10, 64 days vs. 45.5 days, log-rank test, p=0.012). In contrast, PDGF tumor-bearing Rag1^−/-^ mice exhibited similar survival to WT controls (WT *n*=8, Rag1^−^/^−^ *n*=7, 33 days vs. 31 days, log-rank test, p=0.952), Figure 8D. This suggests that the adaptive immune system plays a crucial role in glioblastoma progression in mesenchymal subtype. These combined findings are summarized in a schematic (Figure 8E), which depicts the subtype-specific relationship between CD44-associated migration and immune infiltration, and how these factors differentially influence tumor progression.

## DISCUSSION

Understanding glioma progression and the mechanism driving glioma cell migration is critical for the design of effective therapies. Here we developed high-grade glioma mouse models which capture the transcriptomic and the immune microenvironment changes associated with human proneural and mesenchymal GBMs. Using the mouse models and PDX lines, we defined a mechanistic difference in glioma cell migration which highlights a functional characteristic of GBM molecular subtypes. The migration difference was consistent with changes in cellular adhesion, notably by *CD44*, but not molecular motors such as myosin II motors. This finding points toward an anti-migratory therapy approach targeted against cellular adhesion as opposed to myosin motors. With the negative Phase III clinical trial of the integrin-inhibitor, Cilengitide for GBM (Stupp *et al*., 2014), it is possible that integrins may not be the major adhesion molecules utilized by glioma cells to migrate but instead they could utilize CD44. While anti-CD44 therapies have not been tried in GBM, an anti-CD44 monoclonal antibody therapy (RO5429083, Roche, Basel, Switzerland) has been investigated in Phase I trials in patients with solid tumors and with AML (Menke-van der Houven van Oordt *et al*., 2016; Vey *et al*., 2016). Therefore, an anti-CD44 therapy could provide clinical benefits by slowing glioma migration.

Furthermore, the upregulation of CD44 in mesenchymal tumors is supportive of the existing literature which defines CD44 as a marker of cancer stem cell and EMT (Ponta, Sherman and Herrlich, 2003; Bloushtain-Qimron *et al*., 2008; Polyak and Weinberg, 2009). During EMT, cancer cells take a more mesenchymal migratory phenotype to allow them to migrate through dense ECM and metastasize (Chaffer and Weinberg, 2011). Consistent with prior mechanistic studies demonstrating a direct role for CD44 in glioma cell migration (Klank et al., 2017b), our results associate enhanced migration in mesenchymal glioma cells with increased traction forces due to increased adhesion molecules expression ‘clutches’ such as CD44. Additionally, Anderson et al. demonstrated a functional role for CD44 in glioma cell migration using brain tissue slice assays, in which both CD44 knockout cells and anti-CD44 monoclonal antibody treatment resulted in reduced migration (Anderson, Kelly and Odde, 2024). Similarly, in breast cancer cells, TGF-β-induced EMT is associated with increased traction forces and clutch number (Mekhdjian *et al*., 2017). Interestingly, downregulation of NF1, a negative regulator of Ras, in epithelial breast cancer cells and Schwann cells also induces expression of transcription factors related to EMT (Arima *et al*., 2010). Moreover, in our study, NRAS^G12V^ expression was used to mimic NF1 downregulation and inactivation in mesenchymal GBM (Verhaak *et al*., 2010; Krusche *et al*., 2016). Ras hyperactivation of MAPK pathway is required for EMT but not PI3K activation by Ras (Janda *et al*., 2002). Altogether, our results implicate EMT in enhanced glioma cell migration and force transmission associated with increased molecular clutches through, and suggest upregulation of clutches, either integrins or CD44, as a conserved feature of EMT across a range of cancers.

Despite the faster migration of the NRAS/Mesenchymal cells, the anti-tumoral immune response within the NRAS/Mesenchymal mouse model is able to slow disease progression and improve survival despite enhanced migration relative to the PDGF/Proneural mouse model. Such an anti-tumor response could potentially be used to slow disease progression and improve clinical outcome for GBM patients. In both mouse and human GBM, mesenchymal tumors are immunologically ‘hot’ relative to the immunologically ‘cold’ proneural tumors (Doucette *et al*., 2013; Wang *et al*., 2017). Despite the presence of immune cells within mesenchymal tumors, immune suppression leads to tumors ultimately prevailing against the anti-tumoral immune response. Based on these findings, we propose an immune checkpoint inhibition strategy, in combination with an anti-migratory therapy, targeting mesenchymal GBM but not proneural GBM.

Our study utilizes an integrated, state of the art experimental approach to study GBM progression and model GBM molecular subtypes by switching a single oncogenic driver (NRAS^G12V^ ↔ PDGFB), in an immunocompetent background without the need for genetically engineered mouse strains or further breeding. Using live cell and brain slice imaging, we identify key mechanical differences between mesenchymal and proneural tumor cells, with mesenchymal cells have larger cellular spread area, generate larger forces, and migrate faster. The functional differences were all predicted by a motor-clutch model for cell adhesion and migration where mesenchymal cells have an optimal level of CD44-mediated adhesion (clutches) relative to myosin motors, while proneural cells lack sufficient CD44 to match the myosin motor activity. Despite the faster migration, NRAS/Mesenchymal mice live longer, consistent with the presence of an anti-tumoral immune response that is lacking in PDGF/Proneural mice. This was further validated using immunodeficient mice: NRAS tumors showed worsened survival in Rag1^−/-^ mice, whereas PDGF tumors showed no difference in survival between WT and Rag1^−/-^ animals. Such dynamics are readily captured computationally with little parameter adjustment using a 3-D Brownian dynamics tumor simulator (BDTS). Our results support a model in which mesenchymal GBM exhibits high CD44 expression and strong immune infiltration, whereas proneural GBM shows lower CD44 and minimal immune activity (Figure 8E). Overall, this work establishes an integrated *in vivo* genetic and biophysical modeling framework to connect animal model and human transcriptionally-defined subtypes to fundamental mechanistic understanding, which has the potential to enable a new modeling-centric approach to clinical translation with application in a wide range of cancers (Brubaker and Lauffenburger, 2020). While our model focuses on comparing mesenchymal and proneural GBM subtypes by altering a single oncogenic driver, it doesn’t fully capture the cellular heterogeneity found in human GBM, where different subtypes and cell states often coexist within the same tumor (Patel *et al*., 2014; Neftel *et al*., 2019). That said, this controlled system allows us to isolate and study the behaviors of each subtype in an immunocompetent setting. This approach is valuable for identifying subtype-specific mechanisms and for building predictive models. Future work incorporating additional mutations, lineage tracing, or single-cell analysis could help bridge the gap between this model and the cellular diversity seen in patients.

## AUTHORS CONTRIBUTION

GAS, BLK, BRT, SSR, DAL and DJO contributed to study initiation, conception and design.

GAS, CJL, BCB and DJO contributed to writing the manuscript

GAS, BCB, RL and JMF contributed to developing mouse tumors

GAS, SKR and ALS contributed to the analysis of transcriptomic data.

GAS ran and analyzed the cell migration simulations

GAS and CJL contributed to the acquisition and analysis of glioma cell migration

GAS established tumor lines and performed traction force measurements

GAS, BCB and HBC contributed to imaging and analysis of histological sections

NG, PCR, DM and DJO contributed to the design and implementation of the Brownian Dynamics Tumor Simulator

All authors contributed to the revisions of the manuscript

## ACKNOWLEDGMENTS

The authors would like to thank Drs. Chris Wilke and Clark C. Chen for helpful discussion. This work was supported by National Institutes of Health/National Cancer Institute grants U54 CA210190 to SSR, DAL and DJO, P01CA254849 to DAL and DJO, U54 CA268069 to DJO and U54CA210180 to JNS. DAL acknowledges the American Cancer Society Research Professor grant, the John and Jean Hedberg Brain Tumor Fund, and the Children’s Cancer Research Fund. The authors acknowledge the Minnesota Supercomputing Institute (MSI) and University of Minnesota Genomic Center at the University of Minnesota for providing resources that contributed to the research results reported within this paper. We also acknowledge the Comparative Pathology, Cancer Bioinformatics, and Cytogenomics Shared Resources at the Masonic Cancer Center at the University of Minnesota for services.

## SUPPLEMENTARY FIGURE AND TABLES LEGENDS

**Figure S1. Unsupervised clustering of mouse tumor and healthy brain tissue transcriptomic profiles and pathway enrichment analysis.** A) Heatmap showing expression profile of NRAS and PDGF tumors and healthy mouse brain tissues. Heatmap shows existence of three gene clusters: Tumor-specific cluster, normal brain tissue specific cluster and NRAS tumor-specific cluster. B,C&D) Gene ontology analysis of gene clusters was performed using EnrichR.

**Figure S2. Clustering analysis of mouse and human tumors using gene signatures associated with classical, mesenchymal, and proneural subtypes.** A) Heatmap shows the clustering of human GBM samples using subtype-specific gene signatures. B) Heatmap shows the clustering of mouse tumors using subtype-specific gene signatures. C&D) Quantification of subtype gene signatures within human GBM subtypes and NRAS and PDGF mouse tumors. Solid and dashed lines represent mean and median values respectively. Error bars are S.E.M. +p <0.05, * p <0.01, ** p<0.001, *** p<0.0001, **** p<0.00001.

**Figure S3. Differential expression analysis of cell migration genes.** Volcano plot depicting –log_10_(adj p value) calculated using FDR adjusted Student’s t test versus log_2_(average FC) for individual genes in A) mouse tumors and B) human TCGA-GBM

**Figure S4. Quantification of single cell migration and cell morphology.** A, B, & C) *Ex vivo* migration and morphology data from each individual animal. D, E, & F) Migration and morphology data for each individual mouse primary tumor line in mouse organotypic brain slice. G, H, & I) Migration and morphology data for each individual PDX line in mouse organotypic brain slice.

**Figure S5. BL image sequences showing tumor regression in some rare cases.** All three surviving mice in NRAS cohort showed tumor regression.

**Figure S6. Expression of immune checkpoint genes within mouse tumors.** Expression of immune checkpoint genes were upregulated in NRAS tumors relative to PDGF tumors (B) similar to human mesenchymal and proneural (A).

**Video S1.** Related to Figure 3. GFP-positive NRAS and PDGF tumor cells migrating in tumor bearing brain slices over 10 hours. Green shows tumor cells and magenta shows blood vasculature. Scale bar: 100 µm.

**Video S2.** Related to Figure 4A. GFP-positive NRAS and PDGF primary tumor cells migrating in mouse organotypic brain slice over 16 hours. Green shows tumor cells and magenta shows blood vasculature. Scale bar: 100 µm.

**Video S3.** Related to Figure 4D. Proneural and mesenchymal human PDX GBM cells migrating in mouse organotypic brain slice over 16 hours. Green shows tumor cells stained using green DiO membrane dye and magenta shows blood vasculature. Scale bar: 100 µm.

**Video S4.** Related to Figure 8. Brownian dynamics tumor simulator output showing simulated tumor growth.

**Table S1.** Related to Figure 1. FPKM expression of mouse tumors and healthy brain tissues.

**Table S2.** Related to Figure 1C. List of genes identified in each cluster (MC1, MC2, MC3, HC1, HC2, HC4).

**Table S3.** Related to Figure S2. List of previously published subtype gene signatures (CL, MES, and PN gene signatures) and subset with normalized standard deviation >1 in mouse and human datasets

**Table S4:** Cell migration simulator parameter values.

| Symbol | Parameter | Value |
| --- | --- | --- |
| $N_m$ | Total number of motors | 10,000 |
| $N_c$ | Total number of clutches | 250; 750 |
| $A_{tot}$ | Total possible actin protrusion length | 100 $\mu\text{m}$ |
| $v_p^*$ | Maximum actin polymerization velocity | 200 nm/s |
| $k_{mod}^*$ | Maximum module birth rate | $1 \text{ s}^{-1}$ |
| $k_{cap}$ | Module capping rate | $0.001 \text{ s}^{-1}$ |
| $l_{in}$ | Initial module length | 5 $\mu\text{m}$ |
| $l_{min}$ | Minimum module length | 0.1 $\mu\text{m}$ |
| $\kappa_{cell}$ | Cell spring constant | 10,000 pN/nm |
| $n_{c,cell}$ | Number of cell body clutches | 10; |
| $n_m^*$ | Maximum number of module motors | $0.1 \cdot N_c$ |
| $F_m$ | Motor stall force | 2 pN |
| $v_m^*$ | Unloaded motor velocity | 120 nm/s |
| $n_c^*$ | Maximum number of module clutches | 75; 750 |
| $k_{on}$ | Clutch on-rate | $1 \text{ s}^{-1}$ |
| $k_{off}^*$ | Clutch unloaded off-rate | $0.1 \text{ s}^{-1}$ |
| $\kappa_c$ | Clutch spring constant | 0.8 pN/nm |
| $F_b$ | Characteristic clutch rupture force | 2 pN |
| $\kappa_s$ | Substrate spring constant | Variable |

**Table S5:** Characteristics of patient-derived xenograft (PDX) lines used in this study.

| GBM | Xenograft start | Sex | Molecular subtype<br>(RNAseq) |
| --- | --- | --- | --- |
| GBM16 | 2002 | F | mesenchymal |
| GBM39 | 2003 | M | mesenchymal |
| GBM44 | 2003 | F | mesenchymal |
| GBM64 | 2006 | F | proneural |
| GBM80 | 2007 | M | proneural |
| GBM85 | 2008 | M | proneural |

**Table S6:**
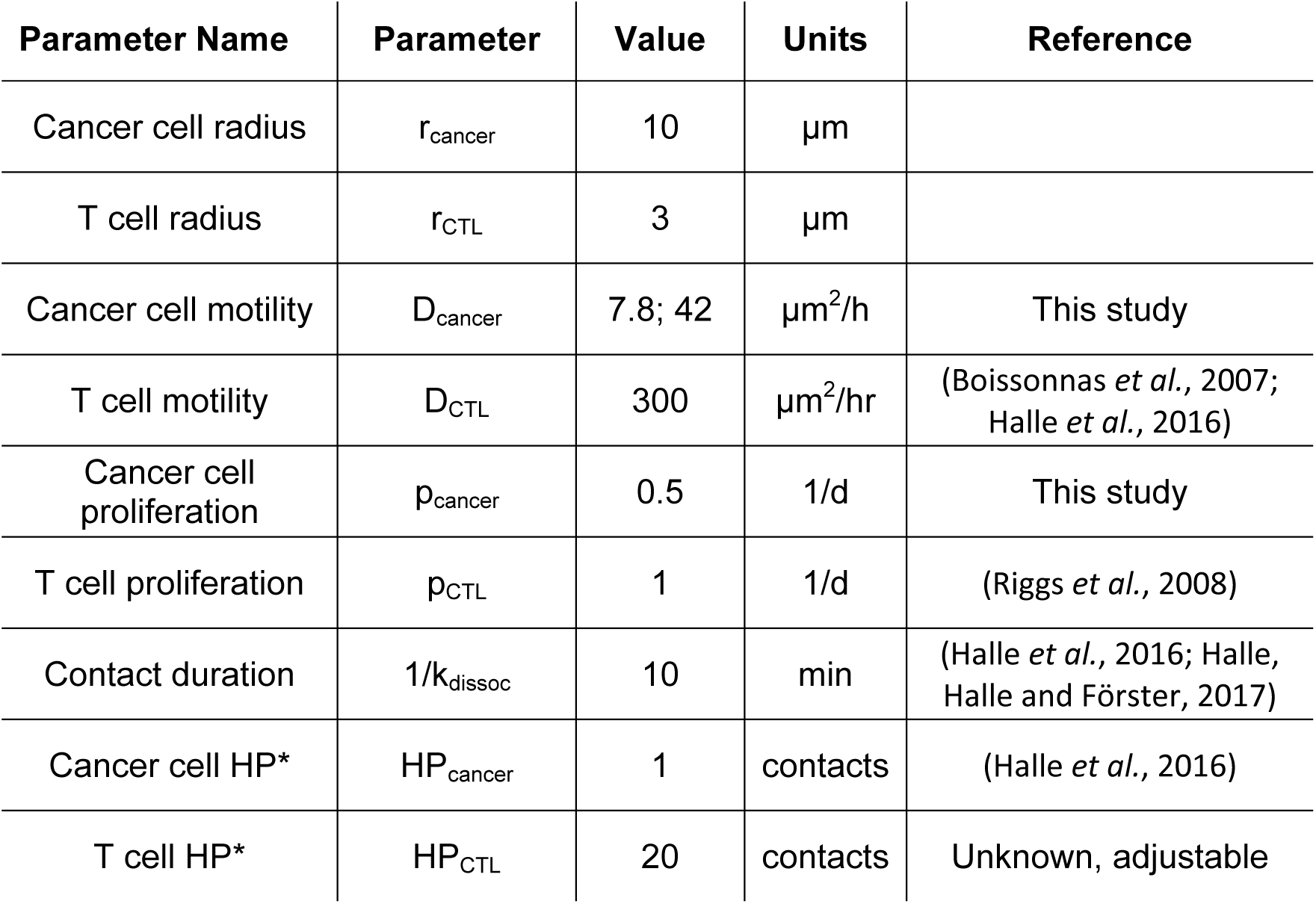
Brownian dynamics tumor simulator parameters values.

**Table S7:** IHC antibodies and reagents used in Figure 1A and Figure 7D.

| IHC | Antibody | Antigen retrieval | Blocking | Detection |
| --- | --- | --- | --- | --- |
| Ki67 | CRM325 | High pH EDTA | Dako protein block | Dako rabbit envision |
| CD3a | A0452 | High pH EDTA | Dako protein block | Biocare rat detection |
| IBA-1 | AB107159 | Low pH citrate | Dako protein block | Biocare goat detection |
| Cleaved Caspase-3 | Asp175 | High pH EDTA | Dako protein block | Dako rabbit envision |
| Granzyme B | D6E9W | Low pH citrate | Vectastain, goat serum | Vectastain ABC-HRP |

