## Supplementary figures and images for "Differential migration mechanics and immune responses of glioblastoma subtypes"

### Figure S1

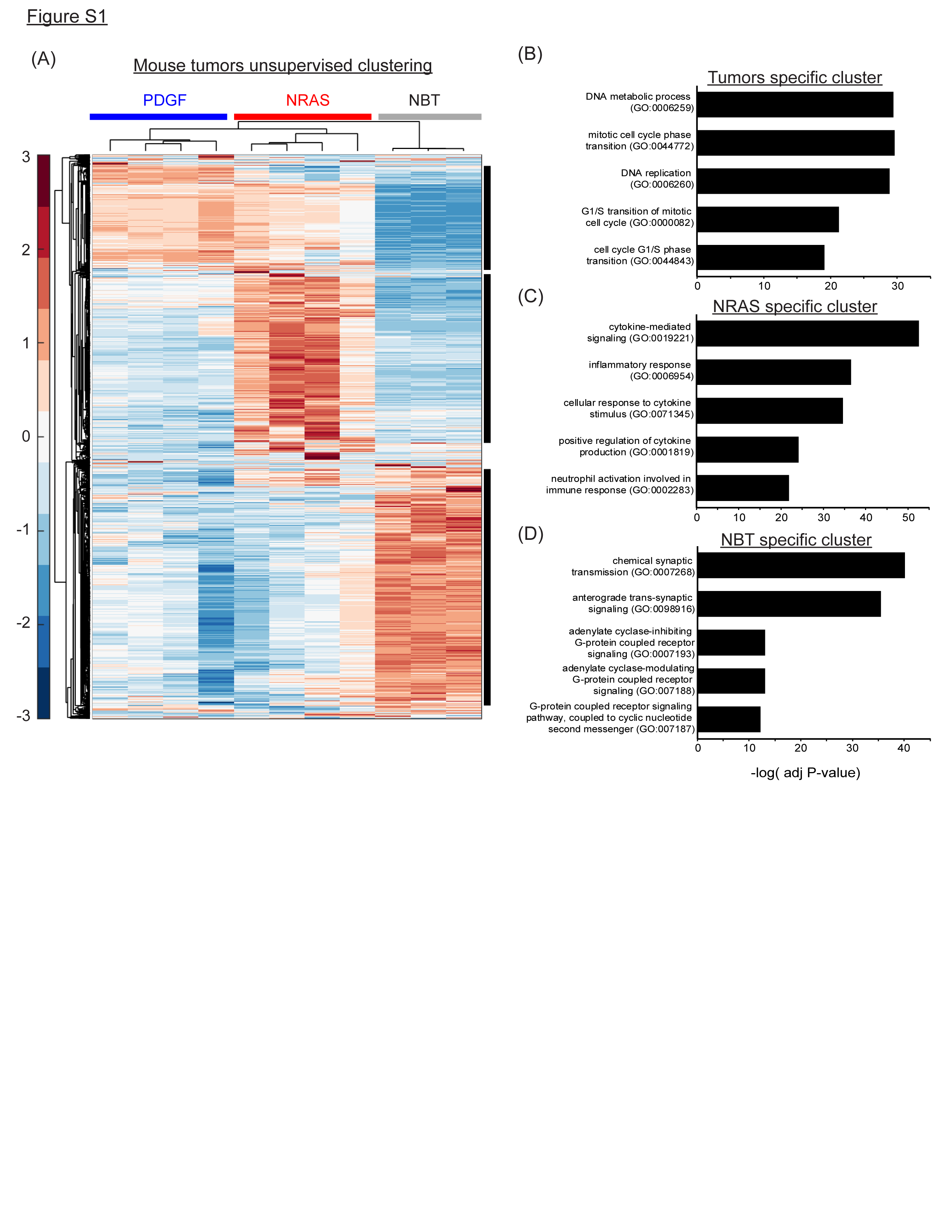

### Figure S2

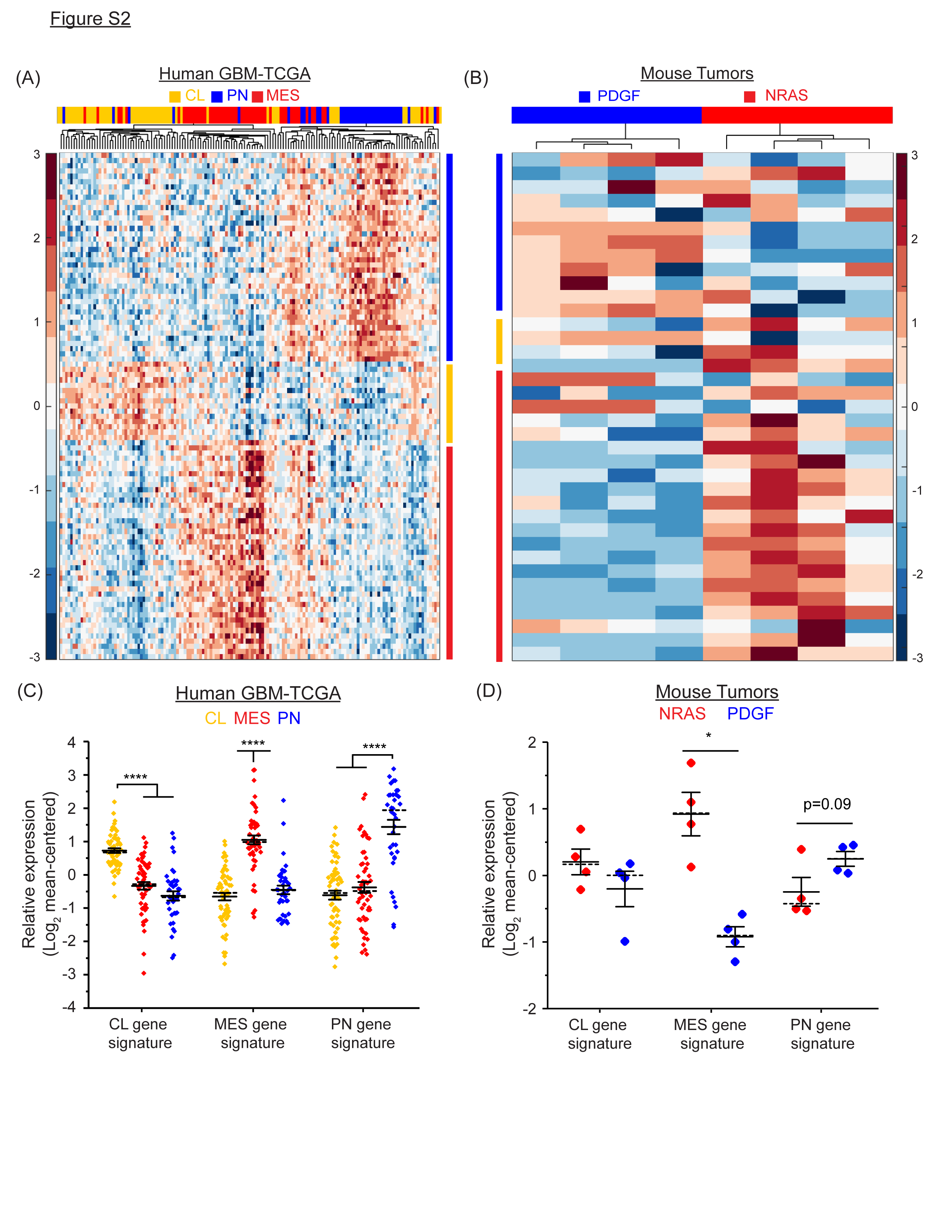

### Figure S3

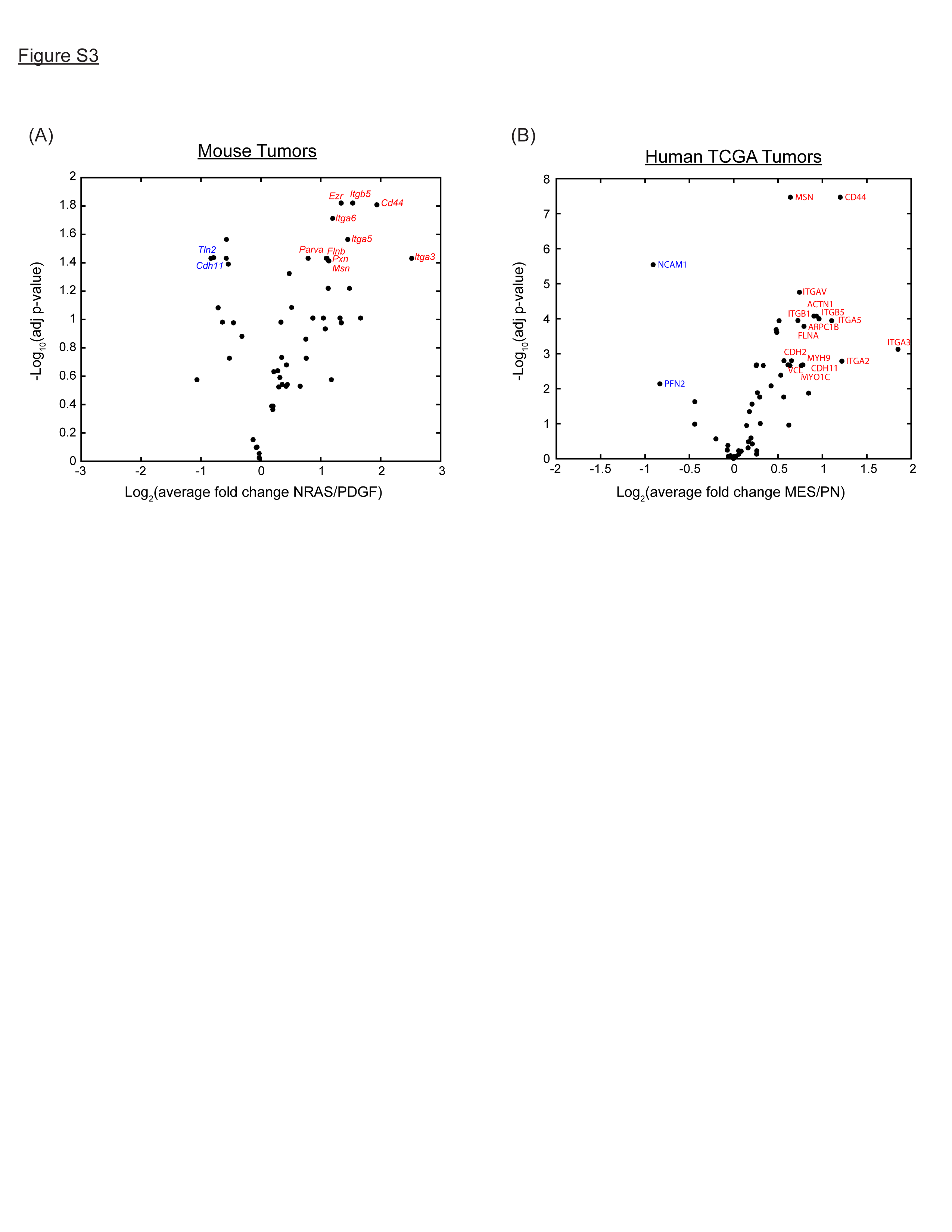

### Figure S4

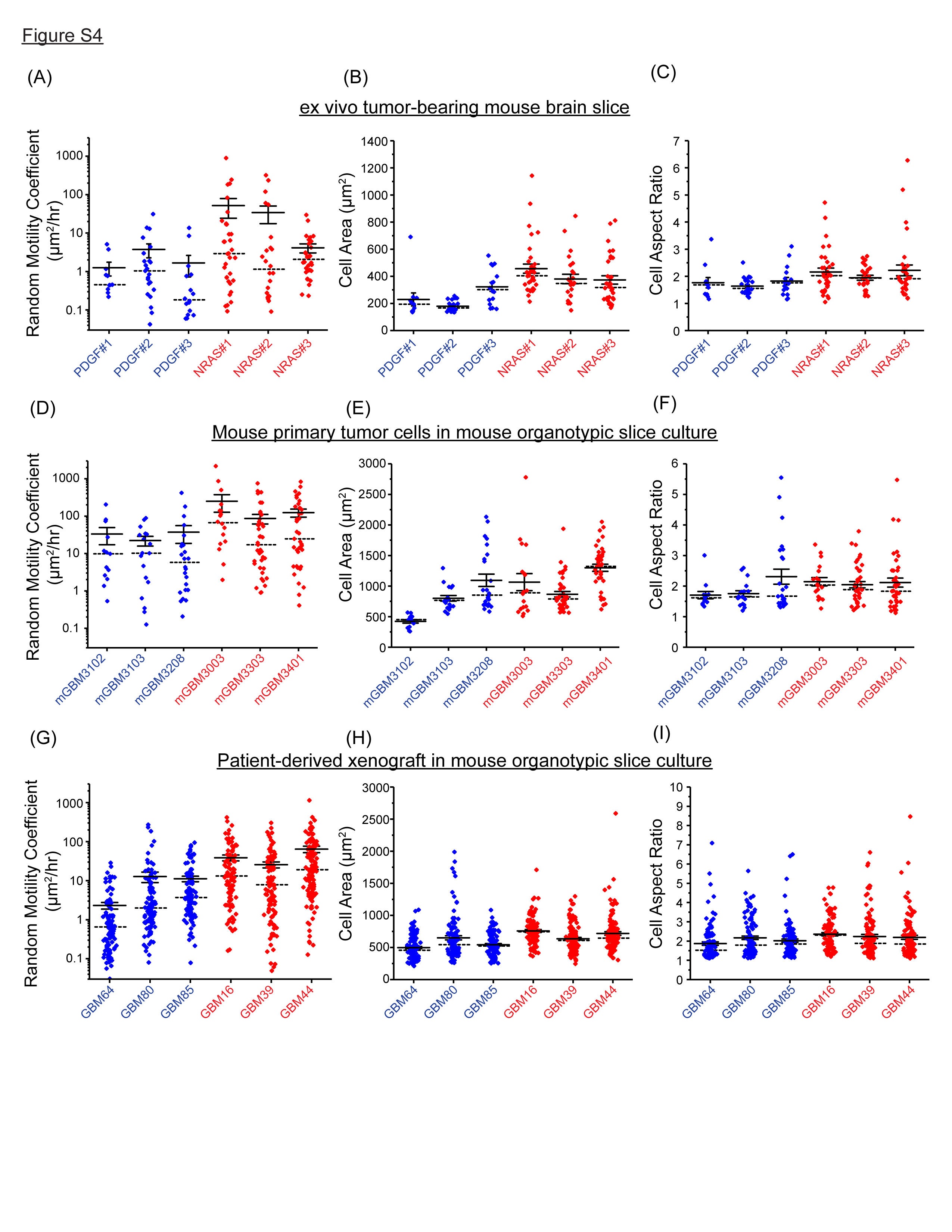

### Figure S5

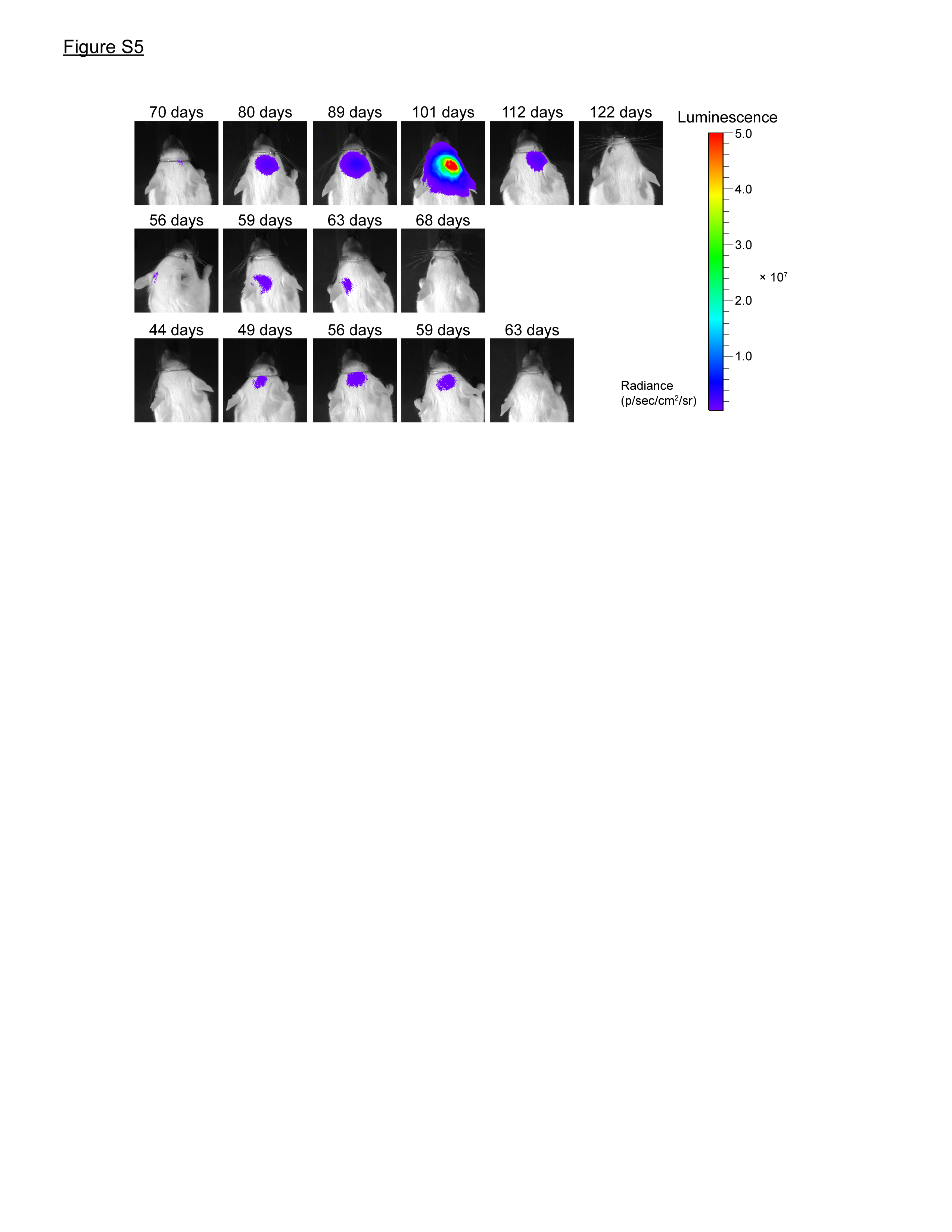

### Figure S6

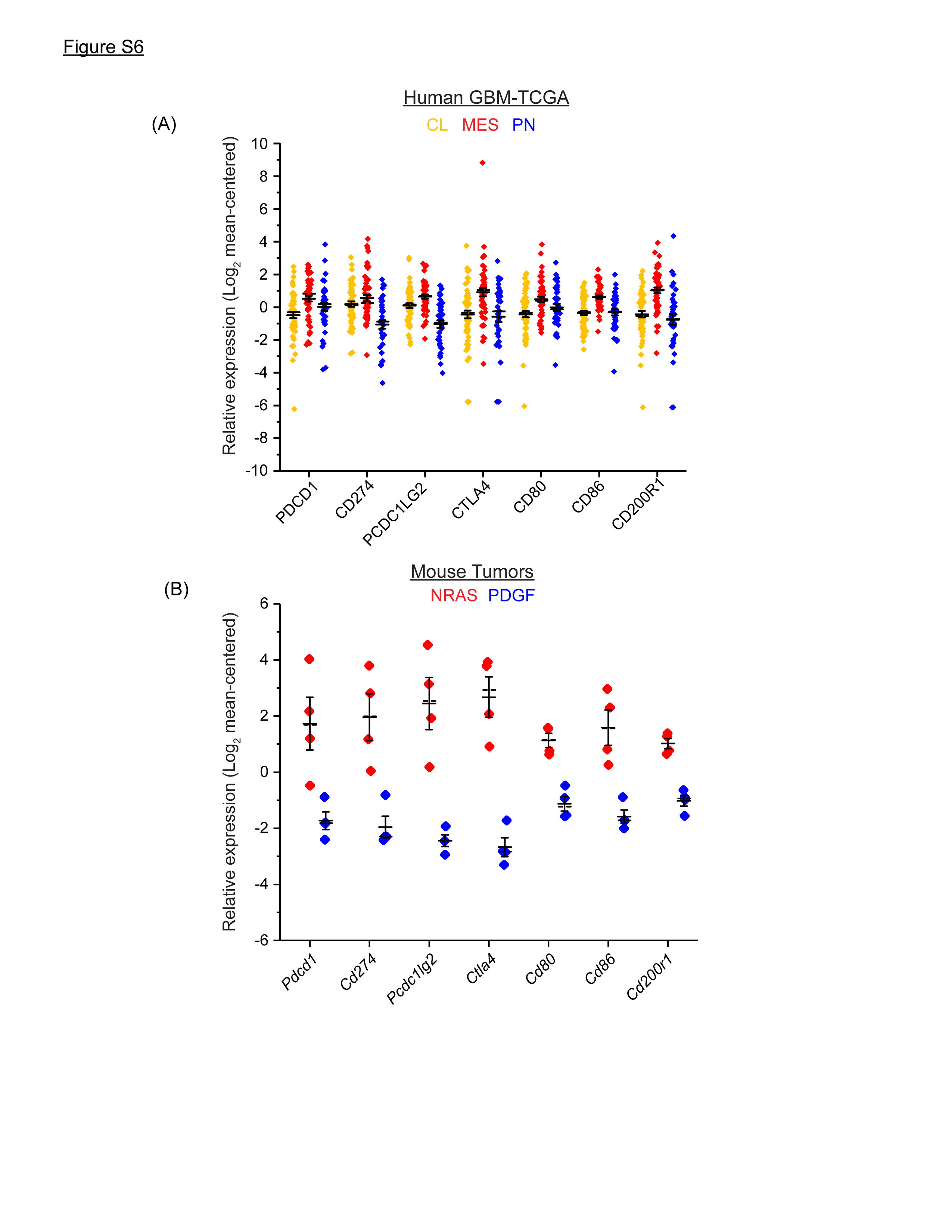
